## Supplemental Information for "Reference-based chemical-genetic interaction profiling to elucidate small molecule mechanism of action in *Mycobacterium tuberculosis*"

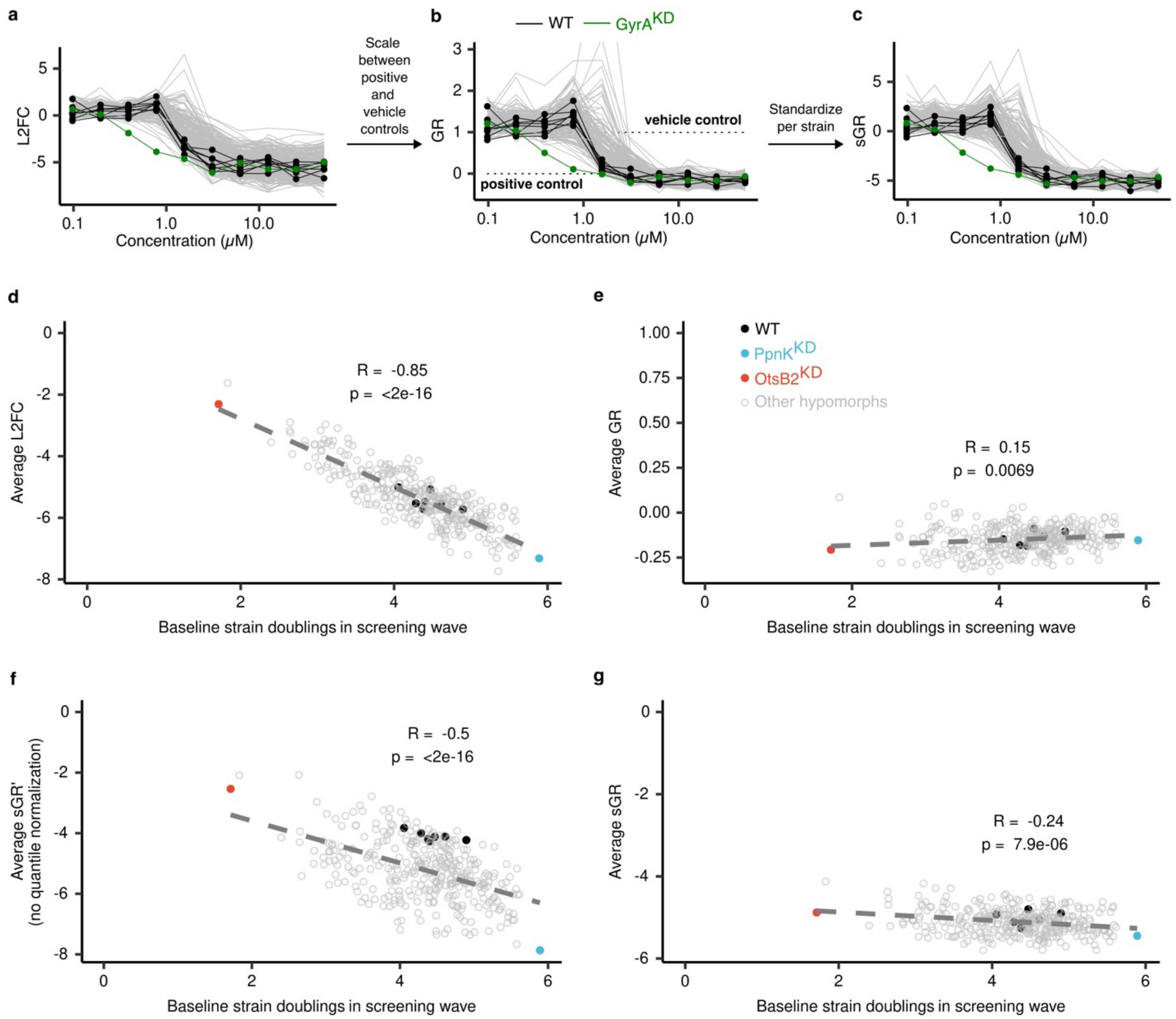


**Extended Data Fig. 1 | Growth scores and strain normalization to reduce dependence on baseline strain growth rates. a-c,** Example growth scores (L2FC, GR, sGR) across different concentrations of the fluoroquinolone nadifloxacin for 7 wildtype H37Rv barcoded strains (black), GyrA hypomorph (green), and 332 other hypomorphs in the pool (gray). Dashed lines represent vehicle (DMSO, GR = 1) and positive (498 nM rifampin, GR = 0) controls used for scaling. **d-g,** Growth scores L2FC, GR, sGR' (= sGR with no strain GR quantile normalization step), and sGR for each strain were averaged over the three highest, fully inhibitory doses of nadifloxacin shown in **a-c** and correlated to each strains' average baseline doublings in the specific screening wave. Black circles are wildtype H37Rv barcoded strains, red circle is the slowest growing strain on average in the screening wave OtsB2, blue circle is the fastest growing strain on average in the screening wave PpnK, and gray open circles are all other hypomorphs. Dark grey dashed line is the best-fit linear regression line and Pearson correlation coefficient and p-value shown. **d,** L2FC is strongly, inversely correlated to strains' baseline doublings, while in **e** GR shows weak-to-no association as expected for doses of compound that completely and equally inhibit the strain pool. **f**, Robust Z scoring reintroduces the inverse correlation (in sGR'), but quantile normalizing strain GR distributions prior to Z scoring, in **g** (sGR), reduces the association. For more detail on mean-variance dependence in GR based on baseline growth and plots for additional example reference set compounds see **Extended Data Fig. 3**.

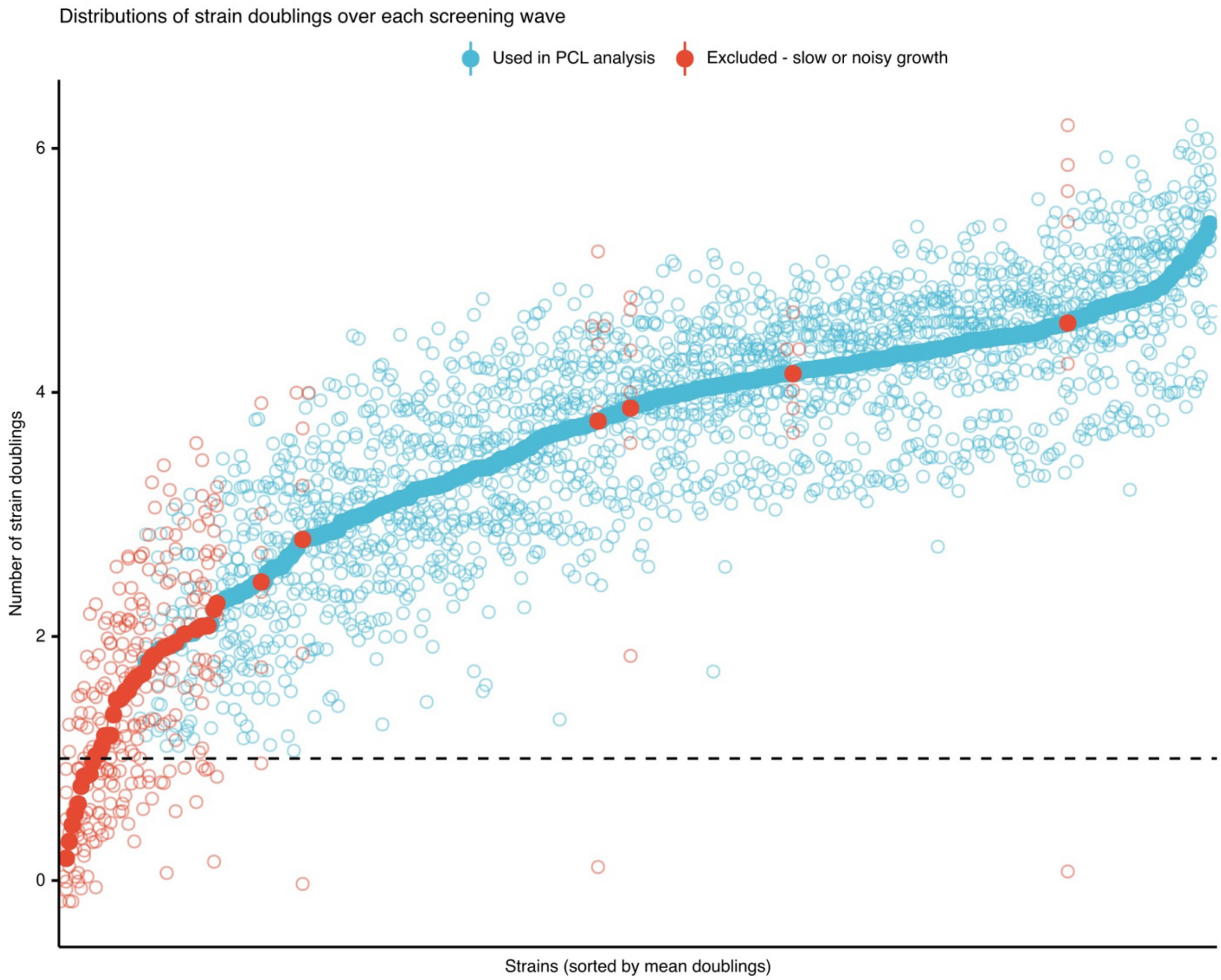


**Extended Data Fig. 2 | Variation in strain doublings/baseline growth rates.** Dot plot of each strain's baseline doublings (open-circle) in the six screening waves and the average across all six waves (filled circle). Black dashed line represents a 1 doubling cutoff for slow growth. Data for 388 strains are shown: 340 (blue) that passed quality control and were used in the PCL analysis (i.e., grew at least 1 doubling in all six screening waves and no conditions with GR >50) and 48 (red) that failed quality control and were excluded (i.e., 45 for slow growth, <1 doubling in at least one screening wave and 3 for noisy growth, conditions with GR >50). 83 strains (not shown) were present in at least one but not all the screening waves and were excluded from the combined PCL analysis. For underlying data and strain identity see **Supplementary Data 2**.

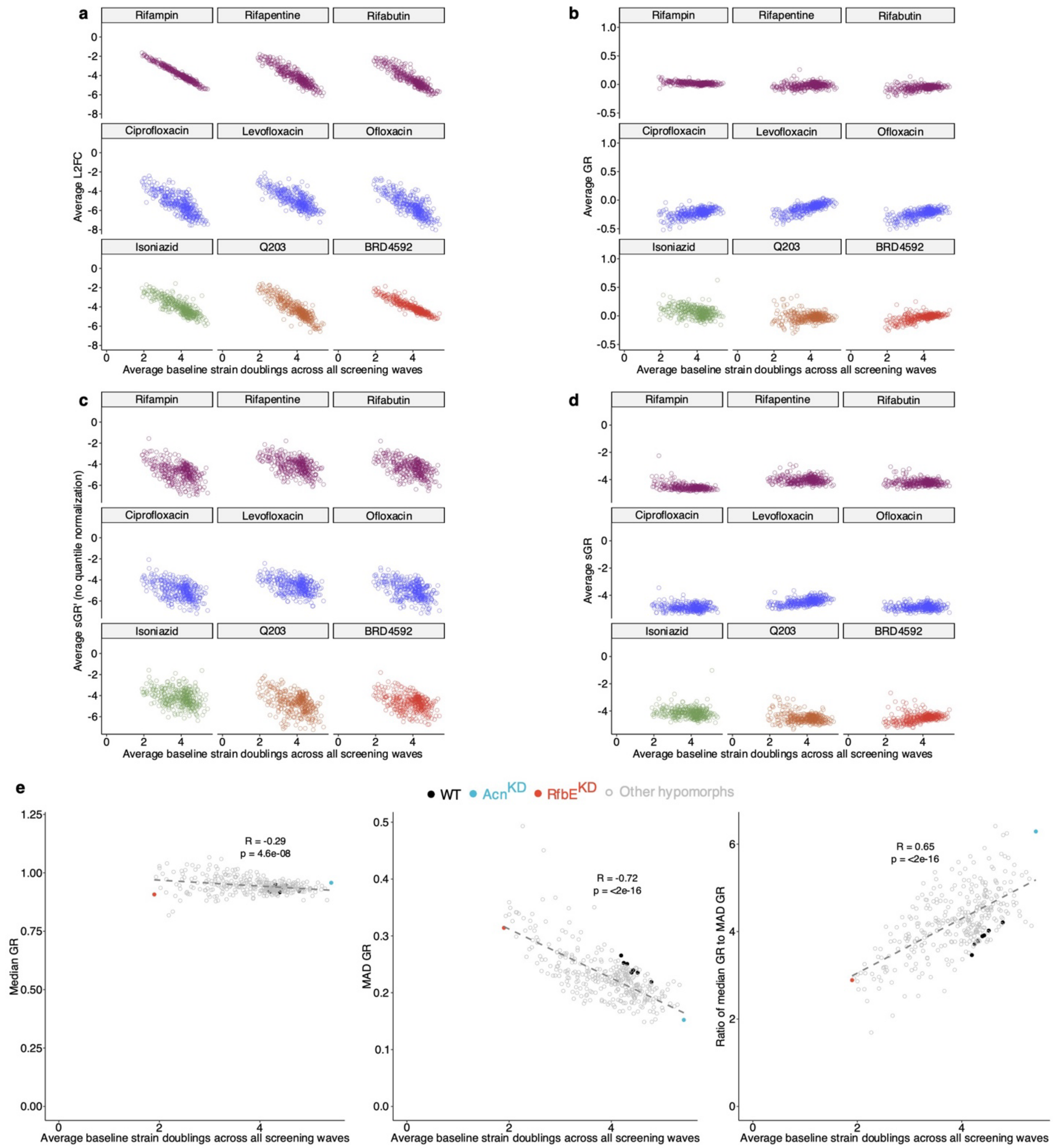


**Extended Data Fig. 3 | Mean-variance dependence in GR based on baseline growth and addressing dependence for hit-calling through quantile normalization.** As in **Extended Data Fig. 1d-g,** growth scores L2FC, GR, sGR' (= sGR with no strain GR quantile normalization step), and sGR for each strain were averaged over fully inhibitory, >4X MIC doses of 9 diverse, exemplary reference set compounds (62 total doses) and correlated to each strains' average doublings (L2FC of onboard, 256 nM or 498 nM rifampin) across the screening waves. Average growth scores for each strain were calculated and plotted separately for each compound (rifamycins in purple, fluoroquinolones in blue, isoniazid in green, Q203 in orange, and BRD4592 in red). **a,** L2FC is strongly, inversely correlated to strains' base doublings while in **b** GR shows weak-to-no association, as expected for doses of compound that completely and equally inhibit the strain pool. **c**, Robust Z scoring reintroduces the inverse correlation (in sGR'), but quantile normalizing strain GR distributions prior to Z scoring, in **d** (sGR), reduces the association. For reference set compounds and doses considered fully inhibitory in **a-d** see **Supplementary Note 3**. **e**, Scatter plots of each strains' average doublings versus their (**left**) median GR, (**middle)** median absolute deviation (MAD) in GR, and (**right)** ratio of median GR to MAD GR over all screened conditions. Black circles are wildtype H37Rv barcoded strains, red circle is the slowest growing strain on average across all screening waves RfbE, blue circle is the fastest growing strain on average across all screening waves Acn, and gray open circles are all other hypomorphs. Dark gray dashed line is the best-fit linear regression line and Pearson correlation coefficient and p-value shown.

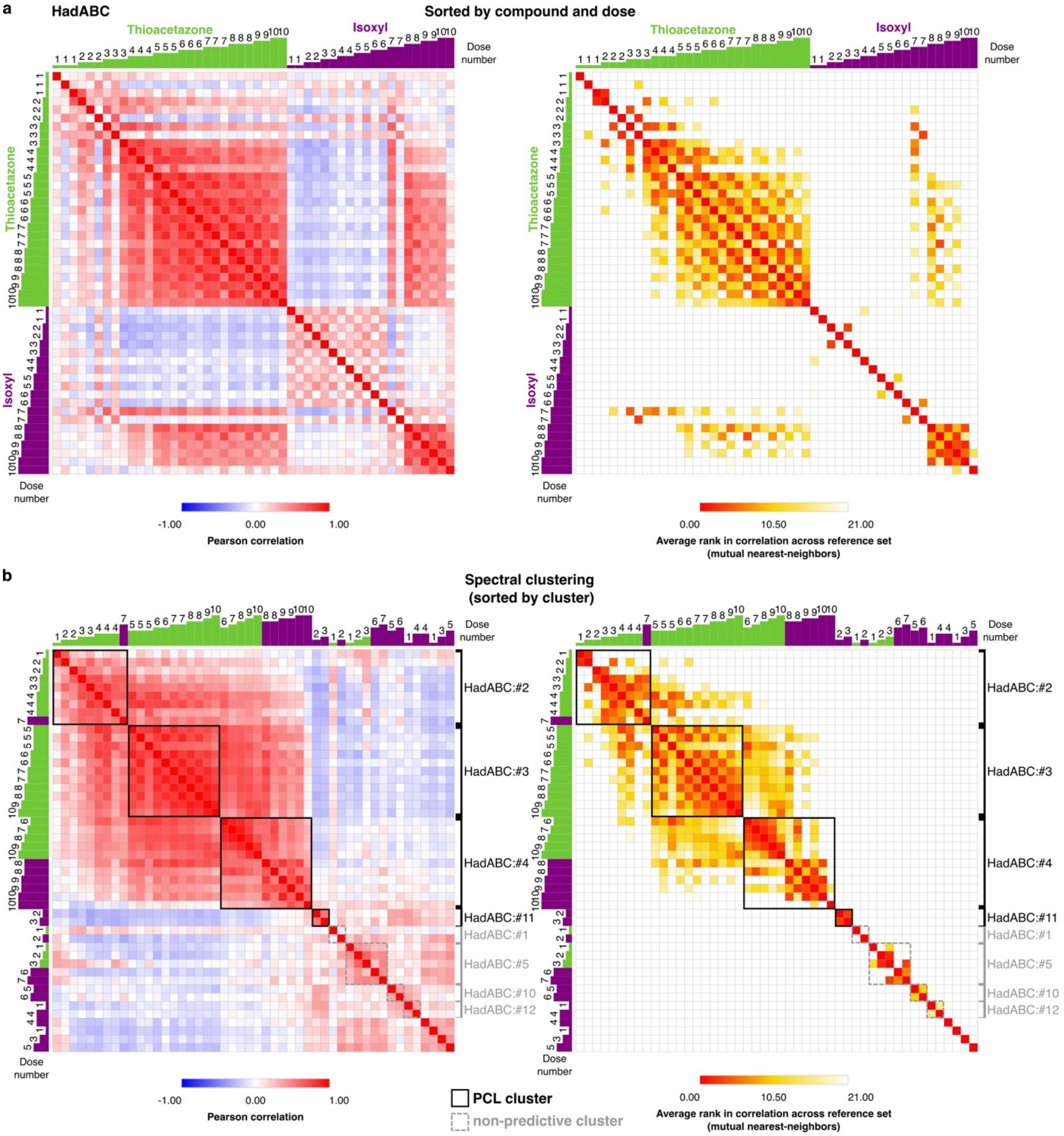


**Extended Data Fig. 4 | Spectral clustering results for example MOA HadABC. a,** Heatmaps of (**left**) Pearson correlation between CGI profiles and (**right**) average rank in correlation across all reference set CGI profiles for the two compounds, thioacetazone (green) and isoxyl (purple), with annotated MOA HadABC in mycolic acid synthesis. Thioacetazone was screened in dose-response in three of the six screening waves and isoxyl was screened in dose-response in two. Dose number (1 = 0.1 µM to 10 = 50 µM in two-fold dilution series) is depicted in green/purple bar plots outside the heatmaps. Heatmaps in **a** are sorted by compound and dose from low-to-high concentration. A threshold of ≤ 20 was applied to the (**right**) average rank of correlation matrix to connect HadABC CGI profiles that were mutual nearest-neighbors and create an adjacency matrix (1 = connected conditions, 0 = disconnected conditions) as input for spectral clustering. For more detail on the steps of spectral clustering, see **Supplementary Note 7**. **b**, as in **a** but heatmaps re-sorted by cluster membership from spectral clustering. PCL clusters and non-predictive clusters are outlined in black and dashed-gray squares respectively, while singleton CGI profiles that did not cluster with others are unlabeled. For HadABC:#4, its PCL similarity score distribution across all reference set CGI profiles and determination of its high-confidence similarity score threshold is shown in **Extended Data Fig. 8a**.

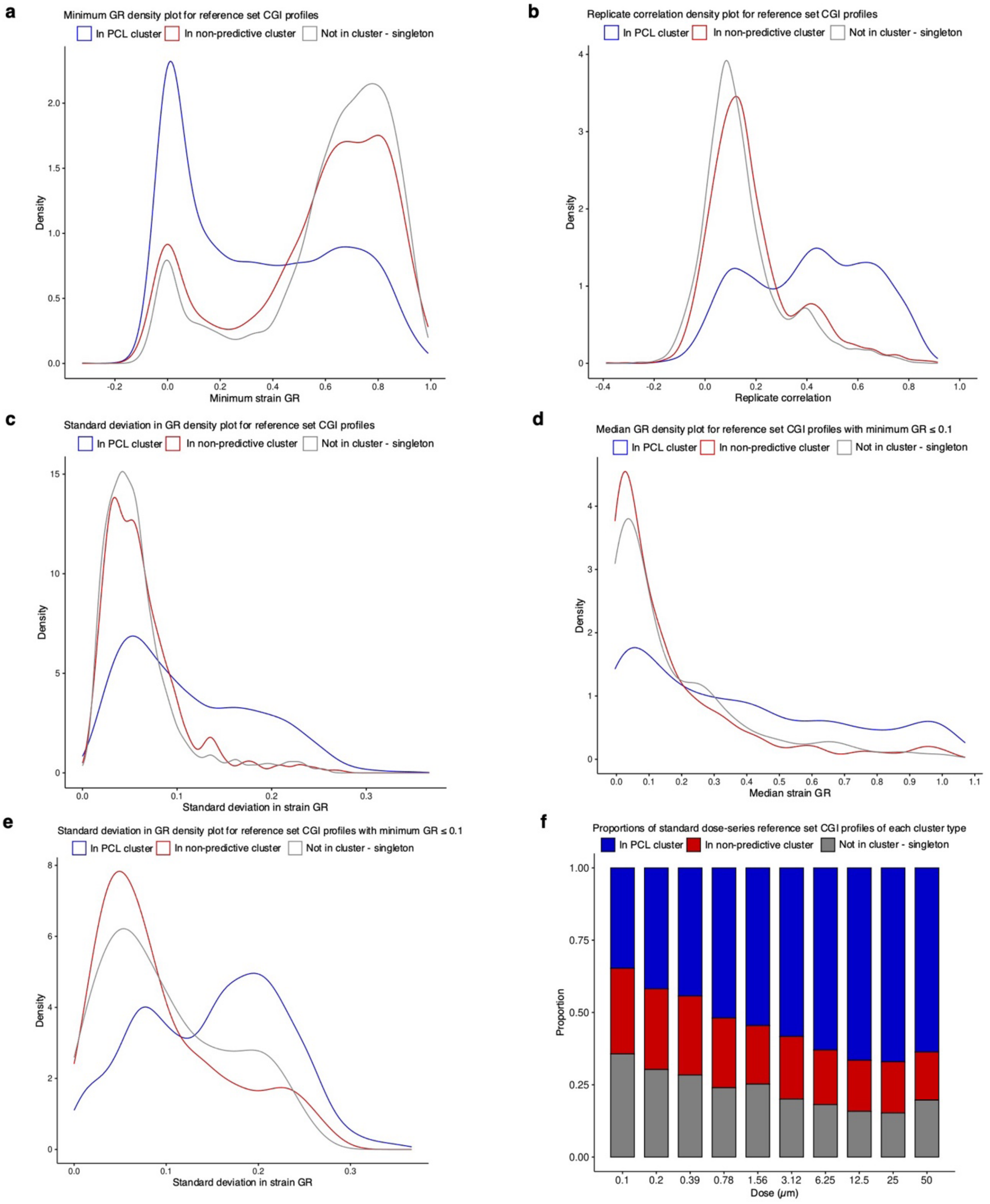


**Extended Data Fig. 5 | Distributions of strain inhibition, replicate correlation, and dose for reference set CGI profiles based on their cluster membership from spectral clustering.** Of the 9,427 CGI profiles in the reference set, 55% (5,134) were members of a PCL cluster (blue), 22% (2,095) were members of a non-predictive cluster (red), and 23% (2,198) were singletons (not in a cluster; gray) in the output of spectral clustering for each MOA (**Supplementary Data 3**). **a**, Density plot of minimum strain GR (maximum strain inhibition) measured for each condition demonstrates that CGI profiles in PCLs tended to have had greater activity with a large proportion showing nearly complete inhibition of at least one strain (minimum strain GR ≤ 0.1). **b** and **c,** Density plots of condition replicate correlation ($n$ = 2 biologically independent experiments) and standard deviation in strain GR for each reference set CGI profile shows conditions that were not in PCLs tended to have poorer replicate reproducibility and lesser variation in strain GR likely due to lack of real, reproducible biological signal (inhibition) and instead strain shuffling being due to random noise. **d** and **e**, Density plots of median strain GR and standard deviation in strain GR for the 2,263 reference set conditions with nearly complete inhibition of at least one strain (minimum strain GR ≤ 0.1) (78% in PCLs, 13% in non-predictive clusters and 9% singletons) shows that active conditions that were not in PCLs tended to have killed most or all strains entirely (median strain GR ≈ 0 and low variance in strain GR) while those in PCLs exhibited wider variation in strain GR capturing meaningful, biological signal for discriminating between MOAs. All 437 reference set compounds were screened twice in two separate screening waves across a standard, 10-point, two-fold dilution series comprising 8,740 "standard dose-series" CGI profiles. **f**, Proportions of "standard dose-series" CGI profiles that were members of a PCL cluster, non-predictive cluster, or not in a cluster. Underlying GR scores shown were strain curve-fit GR values, as described in **Supplementary Note 6**, which reduced dose-to-dose noise; however, using raw GR values did not significantly change the overall observations.

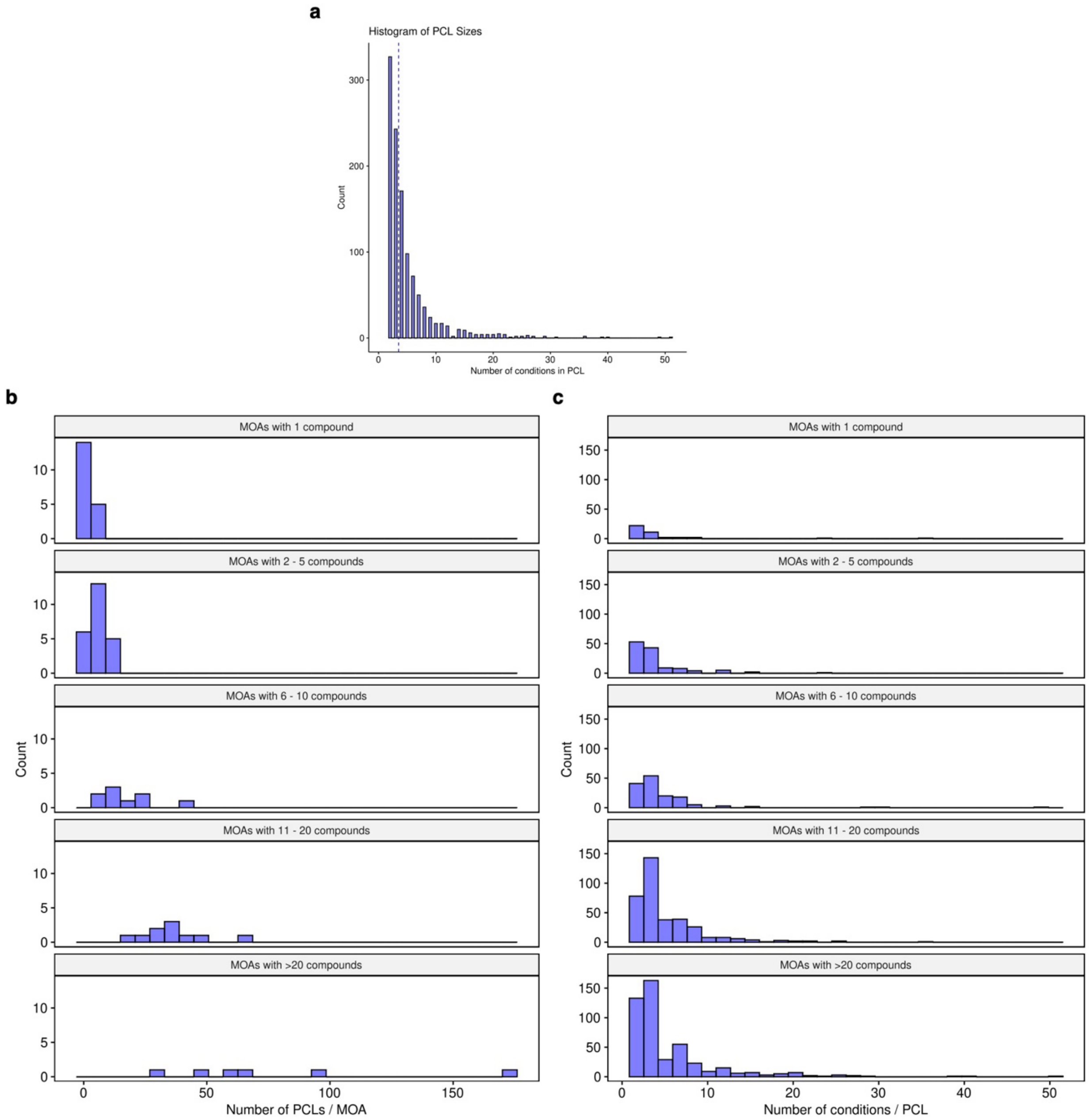


**Extended Data Fig. 6 | Distribution of PCL cluster sizes. a**, Histogram of the number of reference set conditions (CGI profiles) in each of the 1,140 PCL clusters. Blue dashed line indicates the median PCL cluster size of 3.5. **b**, Histograms of the number of PCL clusters constructed per reference MOA for each of the 68 MOAs, grouped by MOA size (i.e., number of reference compounds in the annotated MOA). The number of PCL clusters found via spectral clustering for a given MOA was generally associated with its size. **c**, as in **a**, but histograms grouped by MOA size across the 68 MOAs. Regardless of MOA size, the number of conditions in a PCL cluster varied widely while larger MOAs showed greater enrichment for large PCLs. In both **b** and **c**, 3 single-compound MOAs are excluded for which no PCL clusters were found.

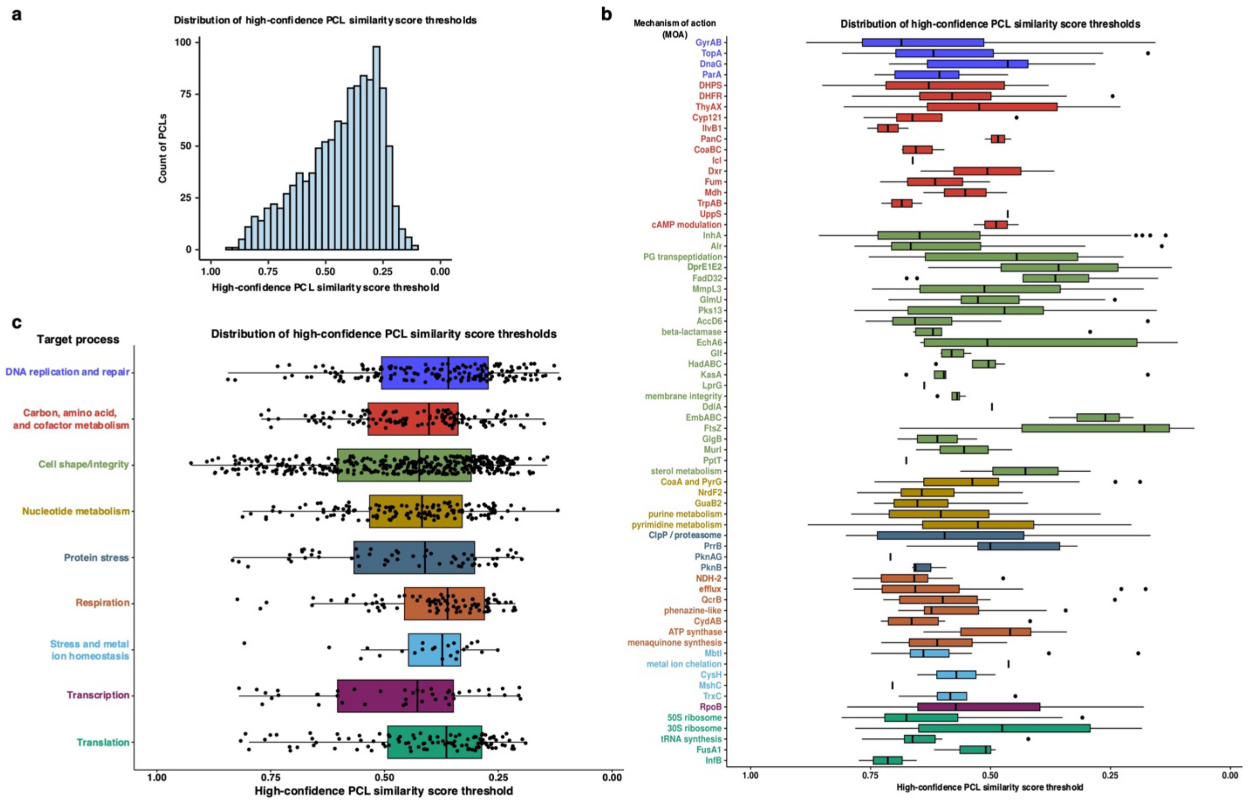


**Extended Data Fig. 7 | Distribution of high-confidence PCL similarity score thresholds. a**, Histogram of the high-confidence PCL similarity score thresholds defined for each of the 1,140 PCL clusters, where PCL similarity score is the median of the Pearson correlation coefficients between a test CGI profile and all CGI profiles in a PCL (excluding self-similarity if the test CGI profile is in the PCL). As depicted in **Fig. 2c**, each PCL's high-confidence threshold was defined as the PCL similarity score of the furthest (least similar) in-MOA, reference set CGI profile before the closest (most similar) out-of-MOA, reference set CGI profile. This corresponds to the minimum PCL similarity score where PCL confidence score = 1 (for examples, see **Extended Data Fig. 8**). PCLs showed a wide range of high-confidence thresholds, owing to the diversity of MOAs represented in the reference set and the different strengths and patterns of strain sensitivities that they elicited (see **Fig. 3c** and **Extended Data Fig. 11b**). Consequently, considering high similarity alone, and not each PCL's unique distribution of PCL similarity scores, when making MOA predictions for reference set compounds performed worse in LOOCV (**Extended Data Table 1**). **b**, As in **a** but boxplot of the high-confidence PCL similarity score thresholds for each PCL cluster grouped and colored by MOA. **c**, As in **a** but jittered boxplot of the high-confidence PCL similarity score thresholds for each PCL cluster grouped and colored by high-level biological target process as in **Fig. 1a** and **Supplementary Data 1**. in-MOA, sharing the same MOA as the PCL conditions; out-of-MOA, distinct MOA from the PCL conditions.

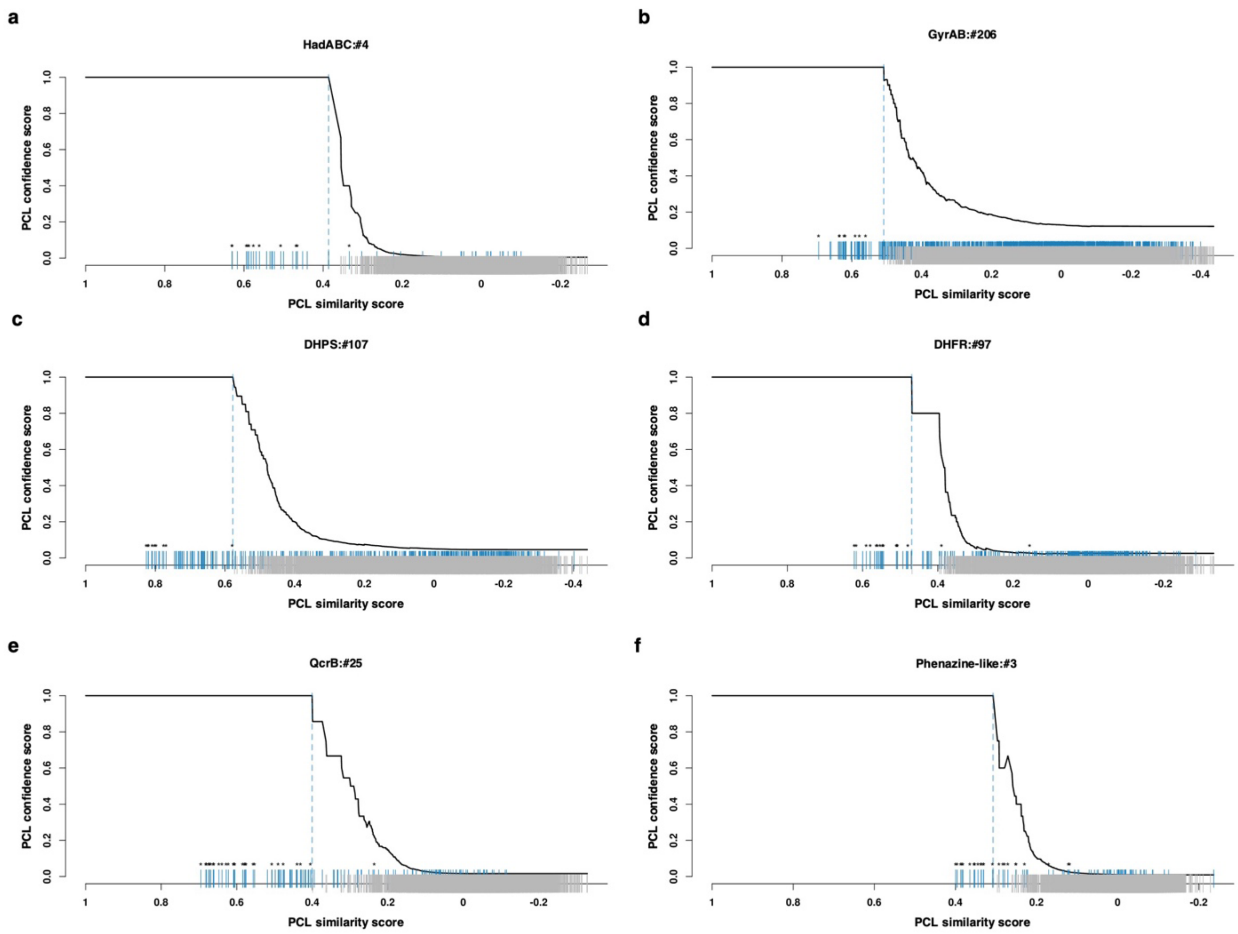


**Extended Data Fig. 8 | High-confidence similarity score thresholds and mappings of PCL similarity score to PCL confidence score for example PCLs.** For 6 representative PCLs, a rug plot of the distribution of similarity scores to a given PCL across all reference set CGI profiles. Overlayed (black line) is the mapping of PCL similarity score to PCL confidence score. As in **Fig. 2c**, blue ticks indicate in-MOA reference set CGI profiles while gray ticks indicate out-of-MOA reference set CGI profiles. Asterisks indicate that the CGI profile is a member of the given PCL cluster. As depicted in **Fig. 2c**, the dashed, light blue line indicates the PCL's defined high-confidence similarity score threshold, i.e., the similarity score of the furthest in-MOA CGI profile before the first out-of-MOA CGI profile. For each observed PCL similarity score $X_i$, PCL confidence score is calculated as the fraction of unique reference set compounds with similarity score $X \geq X_i$ that are in-MOA. In other words, PCL confidence score at a given $X_i$ is the percentage of reference set compounds that are correctly assigned to the PCL's MOA (i.e., precision) if $X_i$ were used as a threshold for which any compound with similarity equal to or above $X_i$ is assigned the PCL's MOA. For a CGI profile from an unknown compound, its confidence score to a particular PCL is estimated by fitting its similarity score to the mapping found empirically using the reference set CGI profiles via linear interpolation and estimates the likelihood of (or proportion of evidence supporting) the unknown compound sharing the PCL's MOA. **a**, PCL of mycolic acid synthesis inhibitors: HadABC:#4, composed of 11 CGI profiles representing 2 unique compounds. **b**, PCL of gyrase inhibitors: GyrAB:#206, composed of 8 CGI profiles representing 8 unique compounds. **c-d**, PCLs of folate biosynthesis inhibitors: DHPS:#107 composed of 9 CGI profiles representing 8 unique compounds, and DHFR:#97 composed of 15 CGI profiles representing 3 unique compounds. **e-f**, PCLs of respiration inhibitors: QcrB:#25 composed of 29 CGI profiles representing 4 unique compounds, and Phenazine-like:#3 composed of 24 CGI profiles representing 4 unique compounds. Strain sensitization of PCLs in **e-f** are shown in **Fig. 3c**. For HadABC:#4, Pearson correlation between HadABC CGI profiles and their average rank of correlation across all reference set CGI profiles is shown in **Extended Data Fig. 4b**.

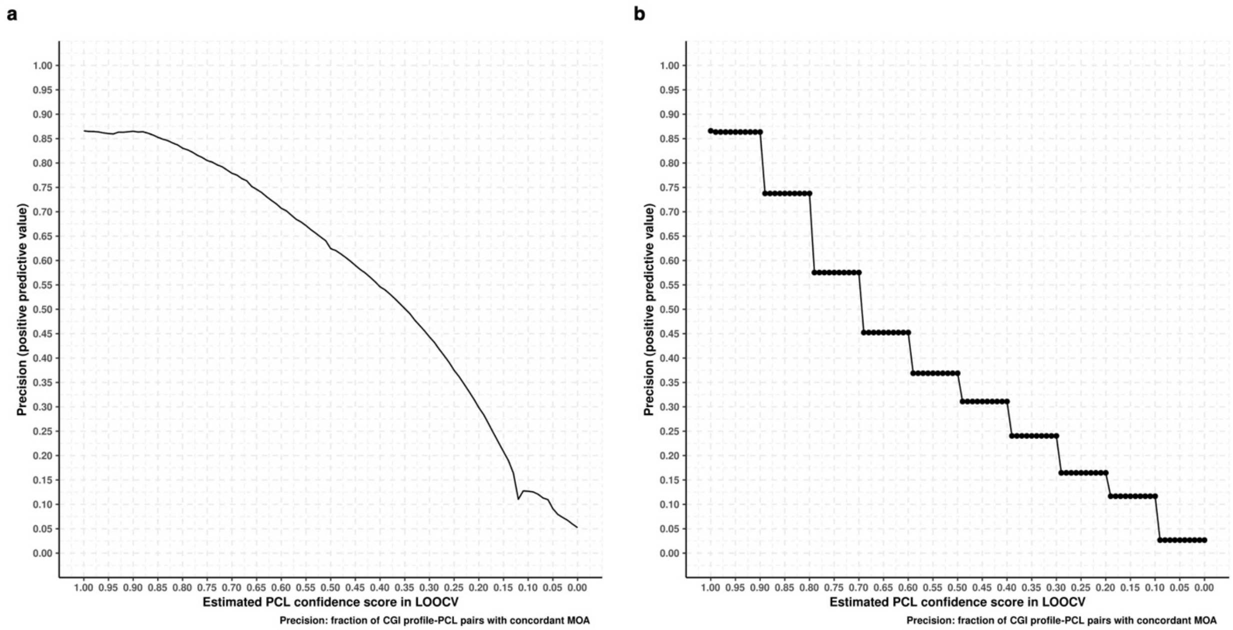


**Extended Data Fig. 9 | Evaluating the predictive value of lower PCL confidence scores.** In LOOCV, each reference set compound was withheld and CGI profiles from the remaining 436 reference set compounds were used in spectral clustering of each MOA to define PCLs, map PCL similarity score to PCL confidence score as in **Extended Data Fig. 8**, and finally estimate the PCL confidence scores for each CGI profile of the withheld reference set compound to every PCL cluster defined in its iteration of LOO. Across all 437 iterations of LOOCV, the median number of PCLs was 1,117 and there were 10,470,531 CGI profile-PCL pairs with varying PCL confidence scores from which we could infer the predictive value of below high-confidence threshold (i.e., PCL confidence score < 1) similarities. Plotted are the proportion of reference set CGI profile-PCL pairs whose MOAs agreed that (**left**) had PCL confidence score ≥ $X_i$ for each 0.01 increment or (**right**) had confidence score of 1, less than 1 to 0.9, less than 0.9 to 0.8, etc. for all bins. As expected, the accuracy of assigning a PCL's MOA to a particular CGI profile declined as the PCL confidence score declined. Matches with PCL confidence score ≥ 0.8 maintained relatively high MOA precision and appear acceptable for consideration when unknown compounds lack a high-confidence prediction and reference-based, MOA information is still of interest. LOOCV, Leave-one-out Cross-validation.

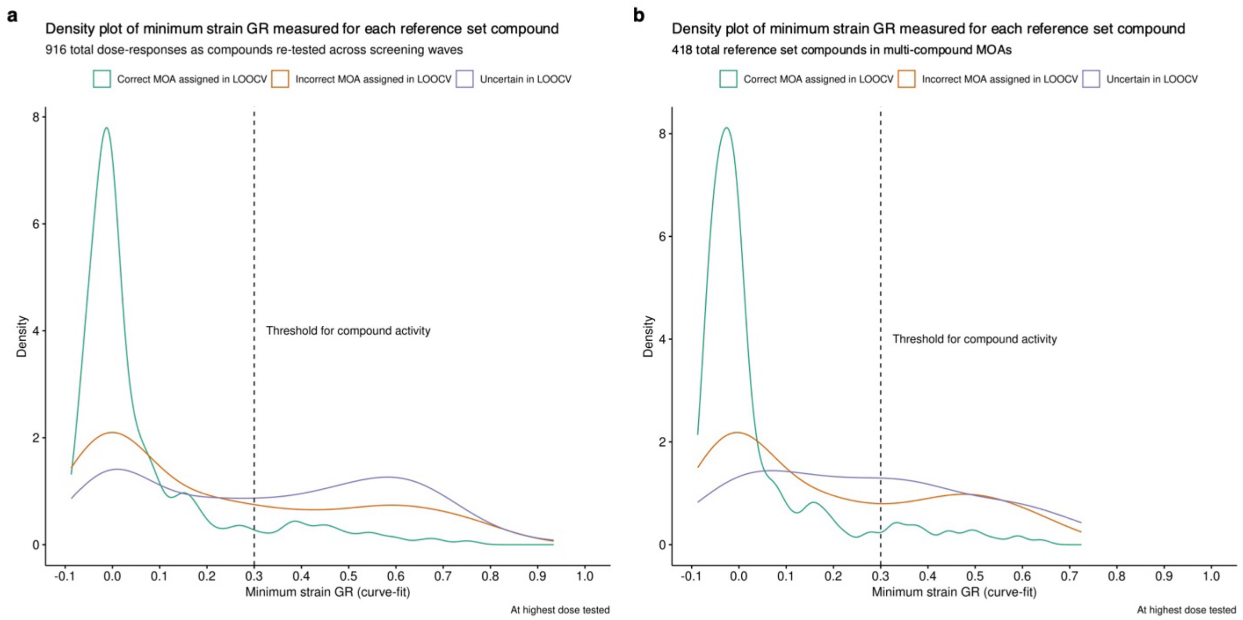


**Extended Data Fig. 10 | Defining compound activity and distributions of minimum strain GR for reference set compounds.** Reference set compounds that were assigned uncertain or incorrect MOA predictions in LOOCV tended to lack clear activity against any strain, whereas compounds that were correctly assigned often inhibited at least one strain completely. Defining a threshold for compound activity aided in identifying which reference and unknown compounds were most likely providing informative biological data, and which would be less trustworthy to make reference-based, MOA predictions for. **a**, Density plots of minimum strain GR achieved at each reference set compound's highest screening concentration in each of the screening waves (916 total dose-series from the 418 reference set compounds that were in multi-compound MOAs). Neglecting for demonstrable compound activity in the screen, the MOA of 501 screening-wave specific instances of reference set compounds would be correctly assigned in LOOCV (green), 187 would be incorrectly assigned (orange), and 228 would be uncertain (purple). **b**, Density plots of minimum strain GR achieved at each reference set compound's highest screening concentration over any one of the screening waves. Neglecting for demonstrable compound activity in any of the screens, 256 reference set compounds would be correctly assigned their MOA in LOOCV (green), while 118 would receive incorrect MOA assignments (orange) and 44 would be uncertain (purple). For both, the black dashed line (minimum strain GR = 0.3) is the threshold we defined for compound activity. Underlying GR scores shown were strain curve-fit GR values, as described in **Supplementary Note 6**, which reduced dose-to-dose noise; however, using raw GR values did not significantly change the overall observations.

**a**

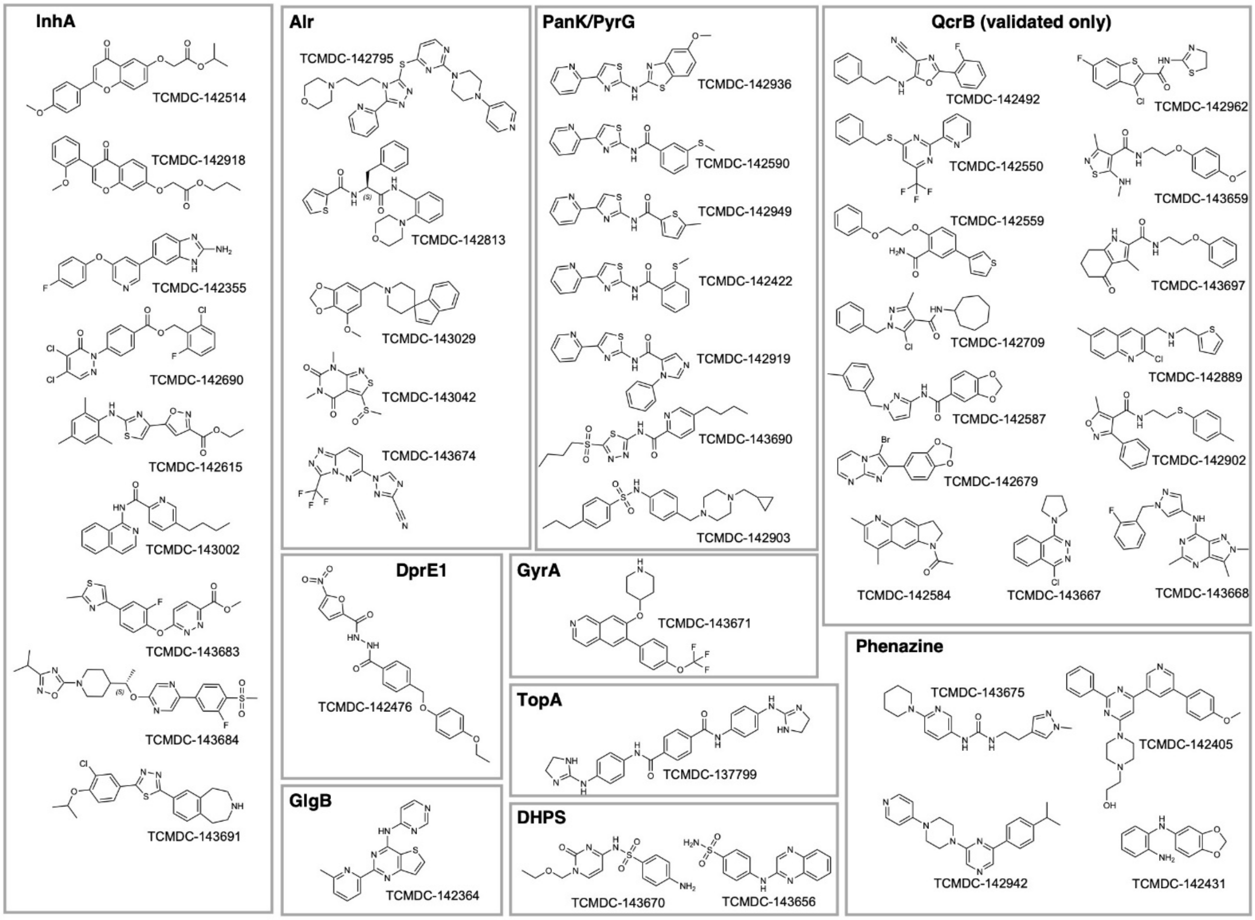


**b**

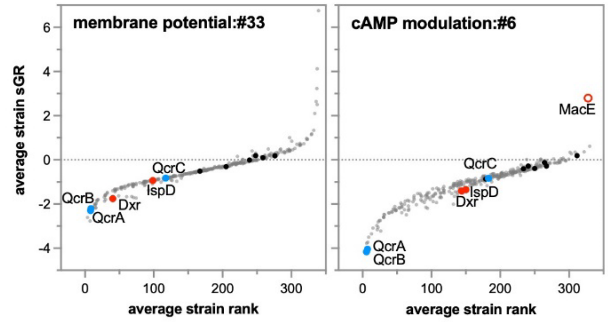


**c**

L2fc: sensitive -5 0 5 resistant

| compound | class | MIC shift: *cydA* | MIC shift: A317T | WT MIC |
|---|---|---|---|---|
| TCMDC-142359 | IMP | -0.5 | 3.0 | 0.1 |
| TCMDC-142376 | | -1.6 | 3.5 | 0.3 |
| TCMDC-142398 | | -2.1 | 1.9 | 0.8 |
| TCMDC-142444 | | -1.2 | 3.6 | 0.5 |
| TCMDC-142511 | | -1.3 | 5.0 | 0.2 |
| TCMDC-142533 | | -0.7 | 4.5 | 0.3 |
| TCMDC-142546 | | -1.4 | 4.2 | 0.7 |
| TCMDC-142557 | | -0.5 | 3.5 | 0.6 |
| TCMDC-142595 | | -0.9 | 4.7 | 0.5 |
| TCMDC-142697 | | -1.3 | 3.3 | 0.5 |
| TCMDC-142785 | | -2.3 | 4.4 | 2.4 |
| TCMDC-142828 | | -0.3 | 5.9 | 0.8 |
| TCMDC-142992 | | -0.5 | 4.9 | 0.4 |
| TCMDC-142490 | PIP | -1.2 | 4.6 | 0.3 |
| TCMDC-142814 | | -1.5 | 2.8 | 5.7 |
| TCMDC-143650 | | -0.6 | 6.2 | 0.2 |
| TCMDC-143652 | | -1.1 | 5.1 | 0.4 |
| TCMDC-143653 | | -1.1 | 3.9 | 1.7 |
| TCMDC-143657 | | -0.6 | 4.0 | 3.2 |
| TCMDC-143698 | | -0.7 | 5.1 | 0.9 |
| TCMDC-142353 | QOA | -3.8 | 1.4 | 1.2 |
| TCMDC-142388 | | -1.4 | 2.6 | 2.0 |
| TCMDC-142461 | | -2.0 | 2.8 | 7.0 |
| TCMDC-142552 | | -1.7 | 2.8 | 14.6 |
| TCMDC-142715 | | -1.0 | 5.9 | 0.8 |
| TCMDC-142430 | QZS | -1.2 | 2.9 | 1.7 |
| TCMDC-142498 | | -2.5 | 3.7 | 0.3 |
| TCMDC-142537 | | -1.2 | 3.1 | 0.2 |
| TCMDC-142368 | TAP | -1.6 | 2.6 | 1.0 |
| TCMDC-142408 | | -1.2 | 2.9 | 3.4 |
| TCMDC-142724 | | -2.2 | 2.9 | 3.4 |
| TCMDC-142980 | PAB | -2.3 | 3.1 | 10.6 |

**control compounds**

| compound | class | *cydA* | A317T | WT MIC |
|---|---|---|---|---|
| Q203 | IMP | -1.1 | 5.8 | 15 nM |
| CFZ | PHEN | -0.8 | -0.3 | 0.6 |

**Extended Data Fig. 11 | Structures of previously unannotated GSK TB set compounds with high-confidence MOA predictions and additional QcrB characterization. a,** Structures of previously unannotated GSK TB set compounds with high-confidence MOA predictions. Only high-confidence predictions (confidence score = 1) are shown. QcrB predictions not supported by MIC shifts in the *cydA*::Tn and $qcrB_{A317T}$ mutants are excluded. **b,** MOA-specific sensitization of cytochrome *bc1* components (QcrA and QcrB) in PCLs formed by non-QcrB-targeting respiration inhibitors. Standardized growth rate (sGR), for each strain, averaged over all compound-dose conditions in the PCL cluster is shown on the y-axis, whereas the average strain rank (based on sGR, 1= most sensitive, 340 = least sensitive) is shown on the x-axis. The PCL membrane potential:#33 is composed of 4 CGI profiles representing consecutive doses of CCCP, and cAMP modulation:#6 is composed of 5 CGI profiles representing consecutive doses of V-12-007960, which has been shown to inhibit adenylate cyclase activity. Cytochrome *bcc* components are highlighted in blue, isoprenoid synthetic enzymes in solid red, and solid black circles are wildtype controls. The MacE hypomorph (open red circle), depleted of an essential adenylate cyclase, shows relative resistance to V-12-007960. **c**, MIC shifts to annotated GSK QcrB inhibitors and controls: Q203 (QcrB inhibitor) and clofazimine in *cydA*::Tn and QcrB A317T mutants, similar to as shown in **Fig. 3**. All 32 of the previously annotated GSK QcrB inhibitors were correctly predicted to target QcrB via PROSPECT and PCL analysis (**Supplementary Data 5**).

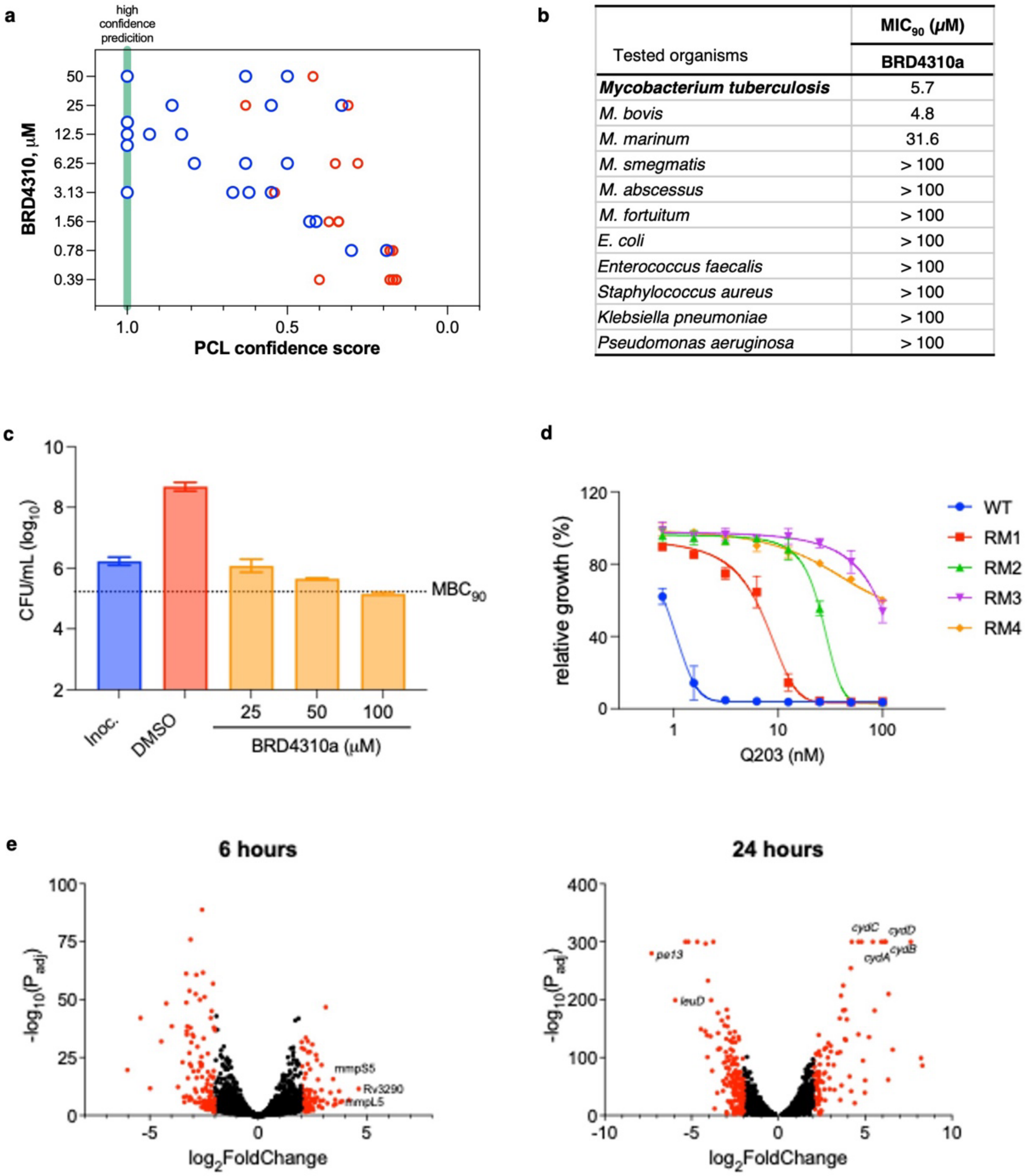


| Tested organisms | MIC$_{90}$ (μM) BRD4310a |
|---|---|
| ***Mycobacterium tuberculosis*** | 5.7 |
| *M. bovis* | 4.8 |
| *M. marinum* | 31.6 |
| *M. smegmatis* | > 100 |
| *M. abscessus* | > 100 |
| *M. fortuitum* | > 100 |
| *E. coli* | > 100 |
| *Enterococcus faecalis* | > 100 |
| *Staphylococcus aureus* | > 100 |
| *Klebsiella pneumoniae* | > 100 |
| *Pseudomonas aeruginosa* | > 100 |

**Extended Data Fig. 12 | QcrB MOA assignment and support for BRD4310 and BRD4310a. a**, Dot plot of PCL confidence scores (X-axis, green line = 1, high confidence) for the top 5 ranking PCLs at each dose of BRD4310, with QcrB PCLs shown in blue and PCLs of other MOAs shown in red. The three high confidence PCL hits are highlighted: QcrB:#25, QcrB:#22, and QcrB:#11. **b**, MIC$_{90}$ of BRD4310a against selected bacteria. **c**, Bactericidal activity of BRD4310a against wild-type H37Rv. Each bar plot shows the mean bacterial burden from biological triplicates and the error bars represent the standard deviation. The dotted line represents 90% bacterial killing compared with the initial inoculum (MBC$_{90}$). All experiments were performed in triplicate and repeated at least once. **d**, Dose–response curves of Q203 against wild-type H37Rv and BRD4310a-resistant mutants. Data are means of triplicates ± SD. **e**, Volcano plot of differential gene expression of wild-type H37Rv exposed to BRD4310a at 6 h and 24 h. Each point represents the average value of one transcript in three replicates.

**Extended Data Table 1 | Accuracy statistics for MOA predictions on validatable, reference compounds in LOOCV.**

| Method | Reference compound activity filter | Total number of reference compounds evaluated | Number of compounds assigned correct MOA | Number of compounds assigned incorrect MOA | Number of uncertain compounds | Precision | Sensitivity | F1 score |
|---|---|---|---|---|---|---|---|---|
| PCL analysis[a] | Active and inactive | 418 | 256 | 118 | 44 | 68% | 61% | 0.65 |
| PCL analysis[a] | Active | 337 | 235 | 77 | 25 | 75% | 70% | 0.72 |
| PCL analysis[a] | Inactive | 81 | 21 | 41 | 19 | 34% | 26% | 0.29 |
| 1-nearest neighbor[b] | Active | 337 | 215 | 122 | N/A | 64% | 64% | 0.64 |
| 1-nearest neighbor[b] | Inactive | 81 | 18 | 63 | N/A | 22% | 22% | 0.22 |
| PCL minus spectral clustering[c] | Active | 337 | 37 | 17 | 283 | 69% | 11% | 0.19 |
| PCL minus individualized similarity score thresholds[d] | Active | 337 | 123 | 34 | 180 | 78% | 36% | 0.50 |
| Most frequent class[e] | Active | 337 | 49 | 288 | N/A | 15% | 15% | 0.15 |
| Random class assignment (average over 1,000 trials)[f] | Active | 337 | 23 | 314 | N/A | 6.8% | 6.8% | 0.07 |

MOA, Mechanism of Action; LOOCV, Leave-one-out cross-validation; CGI, chemical-genetic interaction. Only compounds in multi-compound MOAs can be assigned to the correct mechanism in LOOCV and are considered "validatable". Compounds were defined as active if at least one strain had GR ≤ 0.3 at their highest screened concentration. Micro-averaged precision, sensitivity, and F1 score rounded to two significant figures.

[a] In our PCL analysis method, compounds were assigned an MOA if their similarity score surpassed a PCL's high-confidence similarity score threshold (i.e., PCL confidence score = 1), otherwise they were considered "uncertain". In any case of ties between PCLs of different MOA, the tiebreaker rules selected the PCL for which the highest number of compound-doses (CGI profiles) of the query compound supported the assignment and, if needed, the highest PCL similarity score.

[b] For the 1-nearest neighbor analysis, each compound was queried against the CGI profiles from all other compounds in the reference set and assigned the MOA of the most similar (highest pairwise Pearson correlation) CGI profile over any of its screened doses.

[c] When taking the complete set of CGI profiles from each MOA as one "cluster" per MOA instead of applying spectral clustering, 30 clusters/MOAs were predictive and defined as PCL clusters given that the reference set CGI profile with the highest similarity score to the set was in-MOA, while the other 41 were considered non-predictive as the reference set CGI profiles with the highest similarity score were out-of-MOA (see **Fig. 2c**). Compounds were assigned an MOA if their similarity score surpassed a "PCL" cluster's high-confidence similarity score threshold (i.e., PCL confidence score = 1), otherwise they were considered "uncertain".

[d] Each compound's reference-based MOA prediction was selected as the PCL that it had the highest PCL similarity score to in LOOCV, rather than the highest PCL confidence score. Balancing precision and sensitivity equally, ROC analysis was performed, using the set of PCL similarity scores from each of the reference set compounds and binary labels (1 = agree, 0 = disagree) if the PCL matched the annotated MOA of the compound, to find the optimal similarity score threshold above which a known or unknown compound would be assigned the PCL's MOA with "high-confidence" or otherwise considered "uncertain". Compounds were assigned an MOA if their similarity score surpassed the screen-wide similarity score threshold found in the ROC analysis (*e.g.*, PCL similarity score ≥ 0.741 found for the set of active reference compounds), otherwise they were considered "uncertain". Note that the similarity score threshold found in this approach was more stringent than the high-confidence similarity score thresholds found and defined for most PCLs on

an individualized basis (see **Extended Data Fig. 7**), contributing to significantly lower sensitivity in LOOCV compared to the full PCL analysis.

[e] Trivial model for baseline comparison where each compound's MOA prediction was selected as the most frequent MOA in the reference set: GyrAB.

[f] Trivial model for baseline comparison where each compound's MOA prediction was sampled randomly with replacement from their set (n = 337) of annotated MOAs, maintaining the same imbalanced MOA/class distributions. This was performed and accuracy statistics averaged over 1,000 trials. If the MOA distribution was balanced, the expected average accuracy would be 1 / N for multi-class classification over N classes or 1 / 48 = 2.1% in this case for the active reference compounds.

**Extended Data Table 2 | Accuracy statistics for MOA predictions on annotated GSK set and new PCL predicted, experimentally validated QcrB inhibitors.**

| Method | Total number of compounds evaluated | Number of compounds assigned correct MOA | Number of compounds assigned incorrect MOA | Number of uncertain compounds | Precision | Sensitivity | F1 score |
|---|---|---|---|---|---|---|---|
| PCL analysis[a] | 99 | 72 | 12 | 15 | 86% | 73% | 0.79 |
| Tanimoto similarity[b] | 95 | 36 | 59 | N/A | 38% | 38% | 0.38 |

MOA, Mechanism of Action; CGI, chemical-genetic interaction. Micro-averaged precision, sensitivity, and F1 score rounded to two significant figures. Compounds include the 75 from the GSK set with previous MOA annotation, the 23 from the unannotated GSK set that were predicted by PROSPECT and PCL analysis to target QcrB, and BRD4310. For the annotated GSK, compounds were correctly assigned if the MOA prediction from PCL analysis was concordant with the existing published MOA. For the 23 unannotated GSK predicted to share MOA with reference QcrB inhibitors, the 19 that showed the shifts in MIC between WT, the QcrB resistant mutant strain, and *cydA*::Tn strain consistent with QcrB inhibition were tallied as correct MOA assignments. BRD4310

[a] In our PCL analysis method, compounds were assigned an MOA if their similarity score surpassed a PCL's high-confidence similarity score threshold (i.e., PCL confidence score = 1), otherwise they were considered "uncertain". In any case of ties between PCLs of different MOA, the tiebreaker rules selected the PCL for which the highest number of compound-doses (CGI profiles) of the query compound supported the assignment and, if needed, the highest PCL similarity score.
[b] Trivial model for baseline comparison where each compound's inferred MOA was selected only based on its chemical structure as the reference set compound with the highest Tanimoto similarity to it. The four unannotated GSK compounds that were PCL predicted to target QcrB but failed strain validation are excluded as their exact MOA is still unknown.

**Extended Data Table 3 | Accuracy statistics for PCL-based MOA predictions on reference set in full model (fit to data/training).**

| | Total number of reference set compounds or conditions evaluated | Number of correct MOA assignments | Number of incorrect MOA assignments | Number of uncertain assignments | Precision | Sensitivity | F1 score |
|---|---|---|---|---|---|---|---|
| All reference set compounds (pert_id) | 437 | 411 | 0 | 26 | 100% | 94% | 0.97 |
| All batches of reference set compounds (broad_id) | 505 | 477 | 0 | 28 | 100% | 94% | 0.97 |
| All screening wave instances of batches of reference set compounds (proj_broad_id) | 967 | 850 | 0 | 117 | 100% | 88% | 0.94 |
| All reference set conditions/CGI profiles | 9427 | 3840 | 0 | 5587 | 100% | 41% | 0.58 |

MOA, Mechanism of Action; pert_id, unique identifier for compound based on structure; broad_id, unique identifier for commercial or synthetic lot of a compound; proj_broad_id, unique identifier for an instance of a specific lot of compound screened in a particular screening wave; CGI, chemical-genetic interaction. Compounds or conditions were assigned an MOA if their similarity score surpassed the PCL's high-confidence similarity score threshold (i.e., PCL confidence score = 1), otherwise they were considered "uncertain". The full model refers to the PCLs, high-confidence similarity score thresholds, and mappings of PCL similarity score to PCL confidence score for each PCL that were fit (or learned) using all reference set compounds and applied to all blinded test and unknown compounds. All PCL similarity scores and the true MOA labels for each of the reference set compounds was known when constructing the classifier and thus these statistics on reference set demonstrates the PCL analysis method's fit to and separation of MOAs in the data, not how well to expect the reference-based classification method to generalize to unseen compounds and data. Micro-averaged precision, sensitivity, and F1 score rounded to two significant figures.

### Supplementary Note 1 | Primary analysis of barcode counts and $\log_2$(fold change)

After NGS of the PCR libraries, strain counts for each well were deconvoluted from the undemultiplexed main FASTQ files using the plate (P5 primer), well (P7 primer), and strain barcodes as previously described with the ConCensusMap script[1]. Spike-in control barcode counts in each well were used to $\log_2$-normalize for coverage differences such that median PCR-control barcode counts were equal across wells with different P5 and P7 primers and between different PCR batches (explained in detail below). For every strain, compound, and compound dose combination, $\log_2$(fold change) (L2FC) of spike-in control log-normalized strain counts compared to vehicle control wells were estimated using the ConCensusGLM package as previously described[5].

Each PCR plate had four unique P5 (plate) primers arrayed in quadrant formats (i.e., C2 for P5_1, C3 for P5_2, D2 for P5_3, D3_for P5_4 and so on) and sixty-six P7 (well) primers arrayed in square block format (i.e., C2-C3, D3-D4 for P7_1, …, M22-23, N22-23 for P7_66) which ensured every assay well had a unique P5-P7 primer combination for undemultiplexing strain counts. Using 25 distinct quartets of P5 primers, we were able to include up to 25 assay plates per pooled PCR library with 1,650 total unique P5-P7 primer pairs. The multi-lane architecture of the HiSeq 2500 flow cells enabled the sequencing of many PCR library pools at scale and repeated measurement of each P5-P7 primer pairs' relative efficiencies as estimated via counts of spike-in control barcodes.

Format of Arrayed P5 and P7 Primers for PCR Plates

| | 1 | 2 | 3 | 4 | 5 | 6 | 7 | 8 | 9 | 10 | 11 | 12 | 13 | 14 | 15 | 16 | 17 | 18 | 19 | 20 | 21 | 22 | 23 | 24 |
|---|---|---|---|---|---|---|---|---|---|---|---|---|---|---|---|---|---|---|---|---|---|---|---|---|
| A | | | | | | | | | | | | | | | | | | | | | | | | |
| B | | | | | | | | | | | | | | | | | | | | | | | | |
| C | | P5_1 P7_1 | P5_2 P7_1 | P5_1 P7_2 | P5_2 P7_2 | P5_1 P7_3 | P5_2 P7_3 | P5_1 P7_4 | P5_2 P7_4 | P5_1 P7_5 | P5_2 P7_5 | P5_1 P7_6 | P5_2 P7_6 | P5_1 P7_7 | P5_2 P7_7 | P5_1 P7_8 | P5_2 P7_8 | P5_1 P7_9 | P5_2 P7_9 | P5_1 P7_10 | P5_2 P7_10 | P5_1 P7_11 | P5_2 P7_11 | |
| D | | P5_3 P7_1 | P5_4 P7_1 | P5_3 P7_2 | P5_4 P7_2 | P5_3 P7_3 | P5_4 P7_3 | P5_3 P7_4 | P5_4 P7_4 | P5_3 P7_5 | P5_4 P7_5 | P5_3 P7_6 | P5_4 P7_6 | P5_3 P7_7 | P5_4 P7_7 | P5_3 P7_8 | P5_4 P7_8 | P5_3 P7_9 | P5_4 P7_9 | P5_3 P7_10 | P5_4 P7_10 | P5_3 P7_11 | P5_4 P7_11 | |
| E | | P5_1 P7_12 | P5_2 P7_12 | P5_1 P7_13 | P5_2 P7_13 | P5_1 P7_14 | P5_2 P7_14 | P5_1 P7_15 | P5_2 P7_15 | P5_1 P7_16 | P5_2 P7_16 | P5_1 P7_17 | P5_2 P7_17 | P5_1 P7_18 | P5_2 P7_18 | P5_1 P7_19 | P5_2 P7_19 | P5_1 P7_20 | P5_2 P7_20 | P5_1 P7_21 | P5_2 P7_21 | P5_1 P7_22 | P5_2 P7_22 | |
| F | | P5_3 P7_12 | P5_4 P7_12 | P5_3 P7_13 | P5_4 P7_13 | P5_3 P7_14 | P5_4 P7_14 | P5_3 P7_15 | P5_4 P7_15 | P5_3 P7_16 | P5_4 P7_16 | P5_3 P7_17 | P5_4 P7_17 | P5_3 P7_18 | P5_4 P7_18 | P5_3 P7_19 | P5_4 P7_19 | P5_3 P7_20 | P5_4 P7_20 | P5_3 P7_21 | P5_4 P7_21 | P5_3 P7_22 | P5_4 P7_22 | |
| G | | P5_1 P7_23 | P5_2 P7_23 | P5_1 P7_24 | P5_2 P7_24 | P5_1 P7_25 | P5_2 P7_25 | P5_1 P7_26 | P5_2 P7_26 | P5_1 P7_27 | P5_2 P7_27 | P5_1 P7_28 | P5_2 P7_28 | P5_1 P7_29 | P5_2 P7_29 | P5_1 P7_30 | P5_2 P7_30 | P5_1 P7_31 | P5_2 P7_31 | P5_1 P7_32 | P5_2 P7_32 | P5_1 P7_33 | P5_2 P7_33 | |
| H | | P5_3 P7_23 | P5_4 P7_23 | P5_3 P7_24 | P5_4 P7_24 | P5_3 P7_25 | P5_4 P7_25 | P5_3 P7_26 | P5_4 P7_26 | P5_3 P7_27 | P5_4 P7_27 | P5_3 P7_28 | P5_4 P7_28 | P5_3 P7_29 | P5_4 P7_29 | P5_3 P7_30 | P5_4 P7_30 | P5_3 P7_31 | P5_4 P7_31 | P5_3 P7_32 | P5_4 P7_32 | P5_3 P7_33 | P5_4 P7_33 | |
| I | | P5_1 P7_34 | P5_2 P7_34 | P5_1 P7_35 | P5_2 P7_35 | P5_1 P7_36 | P5_2 P7_36 | P5_1 P7_37 | P5_2 P7_37 | P5_1 P7_38 | P5_2 P7_38 | P5_1 P7_39 | P5_2 P7_39 | P5_1 P7_40 | P5_2 P7_40 | P5_1 P7_41 | P5_2 P7_41 | P5_1 P7_42 | P5_2 P7_42 | P5_1 P7_43 | P5_2 P7_43 | P5_1 P7_44 | P5_2 P7_44 | |
| J | | P5_3 P7_34 | P5_4 P7_34 | P5_3 P7_35 | P5_4 P7_35 | P5_3 P7_36 | P5_4 P7_36 | P5_3 P7_37 | P5_4 P7_37 | P5_3 P7_38 | P5_4 P7_38 | P5_3 P7_39 | P5_4 P7_39 | P5_3 P7_40 | P5_4 P7_40 | P5_3 P7_41 | P5_4 P7_41 | P5_3 P7_42 | P5_4 P7_42 | P5_3 P7_43 | P5_4 P7_43 | P5_3 P7_44 | P5_4 P7_44 | |
| K | | P5_1 P7_45 | P5_2 P7_45 | P5_1 P7_46 | P5_2 P7_46 | P5_1 P7_47 | P5_2 P7_47 | P5_1 P7_48 | P5_2 P7_48 | P5_1 P7_49 | P5_2 P7_49 | P5_1 P7_50 | P5_2 P7_50 | P5_1 P7_51 | P5_2 P7_51 | P5_1 P7_52 | P5_2 P7_52 | P5_1 P7_53 | P5_2 P7_53 | P5_1 P7_54 | P5_2 P7_54 | P5_1 P7_55 | P5_2 P7_55 | |
| L | | P5_3 P7_45 | P5_4 P7_45 | P5_3 P7_46 | P5_4 P7_46 | P5_3 P7_47 | P5_4 P7_47 | P5_3 P7_48 | P5_4 P7_48 | P5_3 P7_49 | P5_4 P7_49 | P5_3 P7_50 | P5_4 P7_50 | P5_3 P7_51 | P5_4 P7_51 | P5_3 P7_52 | P5_4 P7_52 | P5_3 P7_53 | P5_4 P7_53 | P5_3 P7_54 | P5_4 P7_54 | P5_3 P7_55 | P5_4 P7_55 | |
| M | | P5_1 P7_56 | P5_2 P7_56 | P5_1 P7_57 | P5_2 P7_57 | P5_1 P7_58 | P5_2 P7_58 | P5_1 P7_59 | P5_2 P7_59 | P5_1 P7_60 | P5_2 P7_60 | P5_1 P7_61 | P5_2 P7_61 | P5_1 P7_62 | P5_2 P7_62 | P5_1 P7_63 | P5_2 P7_63 | P5_1 P7_64 | P5_2 P7_64 | P5_1 P7_65 | P5_2 P7_65 | P5_1 P7_66 | P5_2 P7_66 | |
| N | | P5_3 P7_56 | P5_4 P7_56 | P5_3 P7_57 | P5_4 P7_57 | P5_3 P7_58 | P5_4 P7_58 | P5_3 P7_59 | P5_4 P7_59 | P5_3 P7_60 | P5_4 P7_60 | P5_3 P7_61 | P5_4 P7_61 | P5_3 P7_62 | P5_4 P7_62 | P5_3 P7_63 | P5_4 P7_63 | P5_3 P7_64 | P5_4 P7_64 | P5_3 P7_65 | P5_4 P7_65 | P5_3 P7_66 | P5_4 P7_66 | |
| O | | | | | | | | | | | | | | | | | | | | | | | | |
| P | | | | | | | | | | | | | | | | | | | | | | | | |

Well count coverage normalization was done in a three-step approach: (1) screening plates from each PCR batch were grouped, the $\log_2$ differences between the median PCR-control count across the PCR batch and the median PCR-control count from each P5 primer (well quadrant) were found as the estimated coverage scaling factors so that all P5 primers would have equal coverage on average, and all strains counts $\log_2$ shifted according to the coverage scaling factor for the P5 primer from that assay well then (2) wells were grouped by P5 primer, the $\log_2$ differences between the median PCR-control count for a P5 primer and the median PCR-control count for each P5-P7 primer pair were found as the next estimated coverage scaling factors so that all P5-P7 primers pairings for a P5 primer would have equal coverage on average, and all strain counts $\log_2$ shifted according to the coverage scaling factor for the P5-P7 primer pair from that assay well and finally (3) screening plates from each PCR batch were grouped, the $\log_2$ differences between the median PCR-control count across the PCR batch and the median PCR-control count from each P7 primer were found as the estimated coverage scaling factors so that all P7 primers would have equal coverage on average, and all strain counts $\log_2$ shifted according to the coverage scaling factor for the P7 primer from that assay well. This spike-in control $\log_2$-normalization technique corrected for systematic, reproducible reductions in coverage (total barcode counts) due to particular primer pairs which would appear downstream as strong compound activity and strain inhibition if unaccounted for.

### Supplementary Note 2 | Growth rate score calculation from $\log_2$(fold change) and strain quality control

Subsequent analyses and data visualization were performed in Matlab and R using the GCTx annotated data matrix format[2]. L2FC was calculated as previously[1]:

$$\text{L2FC} = \log_2\left(\frac{\text{abundance}}{\text{abundance in vehicle}}\right)$$

Dose-dependent compound-induced growth rate (GR) inhibition was calculated from L2FC in reference to the onboard, positive control 256 nM or 498 nM rifampin treated wells which represented complete inhibition of the strain pool at the time of inoculation. Specifically, GR was calculated as:

$$\text{GR} = 2^{\frac{\text{L2FC(rifampin)-L2FC(condition)}}{\text{L2FC(rifampin) - L2FC(vehicle)}}}\text{-1}$$

This is a transformation from the original equation for growth rate, $\text{GR} = 2^{\log_2\left(\frac{x_{\text{trt}}(t)}{x(0)}\right)/\log_2\left(\frac{x_{\text{DMSO}}(t)}{x(0)}\right)}\text{-1}$ in Hafner et al.[3], with strain abundance in onboard, positive control rifampin taken as x(0) to represent counts at time zero. A straightforward derivation from this definition of GR leads to its definition in the main text. While the resulting scale of GR for each strain was approximately between 0 (complete inhibition; rifampin) and 1 (uninhibited growth; vehicle), growth in a competitive, pooled format allowed strains to sometimes have a growth advantage in response to specific conditions and achieve GR > 1 (i.e., abundance and barcode counts greater than vehicle alone). Growth and measurements of GR > 1 is not strictly limited in this assay whereas growth inhibition and GR < 1 is limited more strictly around 0 due to the indestructability of DNA barcodes. GR measurements much greater than 1 were reproducible across biological replicates (i.e., strains had apparent growth advantages across replicate screening plates, multiple doses of the same compounds, multiple compounds of the same MOAs) and we believe reflected real biology (i.e., limited competition and fitness advantage in the *Mtb* hypomorph pool). However, we anticipated that extreme, outlier values of GR >5 could be driven in part by a jackpotting effect during PCR and sequencing where the disproportionate, relative abundance of the strain in the pool was biologically meaningful, but the raw quantity of GR was not biologically reasonable and could interfere with downstream correlation-based analyses. To mitigate for the effect of very large strain GR values when later correlating conditions to one another, any strain GR >5 was re-scaled to GR = 5 + $\log_{10}$(GR).

For each screening wave independently, strains that grew slowly (i.e., L2FC(rifampin) < 1 indicating less than 1 baseline doubling) or unreliably (any conditions with GR > 50) were identified and filtered out from further analysis. Of the 388 strains that were included in the strain pool across all 6 screening waves, 45 strains were filtered out for slow growth in at least one of the screening waves and 3 strains were filtered out for high GR; the 340 strains that passed quality control were used in downstream analysis.

| Strain category | Number of strains | Percent of strains |
|---|---|---|
| Used in PCL analysis | 340 | 87.6% |
| Excluded – slow growth | 45 | 11.6% |
| Excluded – noisy growth | 3 | 0.8% |
| Excluded – not in all screening waves | 83 | - |

Strains were excluded as slow growers if they achieved <1 doubling in one of the six screening waves (see **Supplementary Data 2** for list). MurD, Dfp, and H37RvBC02 were excluded as noisy growers due to many conditions with GR>50.

### Supplementary Note 3 | Standardized growth rate score calculation from GR

Each strain's condition GR distribution was first quantile normalized then robust z-scored using the publicly available CmapM Matlab package to calculate standardized growth rate (sGR), a metric of specific strain sensitivity or resistance relative to the median condition in the entire dataset[4,5]. For each condition and strain, where median(GR) and MAD(GR) are each approximately equal for all strains after quantile normalization:

$$sGR_{treatment,strain} = \frac{GR_{treatment,strain} - median(GR_{strain})}{1.4826 \times MAD(GR_{strain})}$$

GR quantile normalization, over all 340 strains, was performed using the quantilenorm Matlab function taking the median of the ranked values as described in Subramanian et al., in which the researchers similarly standardized expression profiles across cell lines prior to generating differential gene expression[5]. Standardizing the measured growth inhibition across strains according to this protocol served to mitigate mean-variance dependence in GR dependent on baseline strain growth rate which we observed would bias hit-calling and the rank order of strain inhibition to non-specific, completely inhibitory conditions towards how generally fit or quickly strains grew and not due to target-specific or systems biology-specific strain inhibition when challenged with a particular dose of a chemical inhibitor (**Extended Data Fig. 1d-f, Extended Data Fig. 3**). The standardized metric of growth inhibition sGR was on similar scale and units for all strains such that the mean sGR across all conditions is 0 and standard deviation is 1, conditions with sGR much less than 0 indicated extreme strain inhibition relative to the bulk or average of all conditions screened, sGR much greater than 0 indicated extreme strain growth (or resistance) relative to the bulk of all screened conditions, and allowed for inference of each compound's or condition's biological mechanism from the rank order and magnitude of the sGR CGI profile with very little to any dependence on strains' differential growth rates. Quantile normalization and other rank-based and variance-stabilizing transformations have been similarly and commonly used across analysis of microarray and bulk and single-cell RNA-Seq datasets due to mean-variance-dependencies frequently arising from count data (in this case deep sequencing of strain DNA barcodes) that would otherwise make comparisons across cells lines (or strains) inaccurate[6-11]. Each condition's sGR profile (CGI profile) was the basic unit for further analysis making reference-based MOA predictions.

After the calculation of all robust z-score sGR values, CGI profile data for all reference set conditions were visualized using UMAP[12], implemented by the R package UMAP (v 0.2.7.0). Pearson distances between every pair of conditions were calculated as $\sqrt{(1 - \text{Pearson Correlation})}$. The UMAP embedding was performed with the seed set for the random_state and transform_state to 123, number of nearest neighbors for the initial embedding set to 20, and all other default parameters used.

Reference set compounds and doses considered fully inhibitory and included in **Extended Data Fig. 3a-d:**

| Reference set compound | Pert ID | MOA | Number of doses considered fully inhibitory | Dose range considered fully inhibitory | Approximate *Mtb* MIC |
|---|---|---|---|---|---|
| Rifampin | BRD-K01507359 | RpoB | 14 | 0.0244 – 3.13 | 0.006 |
| Rifapentine | BRD-A06889673 | RpoB | 6 | 1.56 – 50 | 0.39 |
| Rifabutin | BRD-K43962705 | RpoB | 13 | 0.0975 – 50 | 0.006 |
| Ciprofloxacin | BRD-K04804440 | GyrAB | 4 | 6.25 – 50 | 1.51 |
| Levofloxacin | BRD-K09471561 | GyrAB | 5 | 3.13 – 50 | 0.69 |
| Ofloxacin | BRD-A24228527 | GyrAB | 4 | 6.25 – 50 | 1.38 |
| Isoniazid | BRD-K87202646 | InhA | 4 | 6.25 – 50 | 0.36 |
| Q203 | BRD-K59853741 | QcrB | 10 | 0.0061 – 3.13 | 0.01 |
| BRD4592 | BRD-K99844592 | TrpAB | 2 | 25 – 50 | 6.25 |

Concentrations shown are micromolar, most rounded to three significant figures. Expected *Mtb* MIC approximated based on published reports and in-house, absorbance-based MIC testing of H37Rv strain.

### Supplementary Note 4 | Replicate correlation

For a descriptive metric of the data quality measured from each condition (screened in at least duplicate), replicate correlation (replicate reproducibility) was calculated as the average Spearman correlation between biological replicates of each condition on or across different assay plates. Specifically, for each well in a 384-well assay plate, strain counts were log2 transformed, robust z-scored to their median counts in the 12 onboard vehicle control (DMSO) wells on each plate, correlated to replicates of that condition in other wells or plates, and the average Spearman rank correlation across replicates taken as the overall replicate reproducibility of the condition in the screen. Any conditions with only one replicate (due to technical error) were assigned null (NAN) replicate correlation values. Robust z-scoring and Spearman correlation calculation was performed using Matlab CmapM library functionality[5].

### Supplementary Note 5 | Combining datasets across screening waves

All primary data analysis (strain barcode counts, L2FC, GR, sGR, and replicate correlation) was performed independently for each screening wave as their own screening datasets. These datasets were combined using GCTx format for further large-scale PCL analysis to include all conditions across the 6 screening waves and the 340 strains that passed quality-control in all waves[2]. For reference set compounds which were assayed in dose-response in at least two screening waves (one earlier against a ~400 strain pool and one later against a ~460 strain pool), the combined dataset had biological replicates (sGR CGI profiles from each individual screening wave) for each condition (compound-dose combination) which both aided in the reference-based PCL analysis and served as internal controls on the reproducibility of PROSPECT.

### Supplementary Note 6 | GR curve-fitting and estimating compound activity

To estimate the activity level of each compound and dose of compound in PROSPECT, the drda R package [13] was used to find the best dose-response curve-fit for each strain's measured GR across a compound's dose series where GR of 1 represents no inhibition (vehicle) and GR of 0 represents full inhibition (256 nM or 498 nM rifampin). Quantile normalized GR was again used to mitigate the effects of strain-to-strain mean-variance dependency that could systematically bias the Hill slopes and upper and lower bounds of each strain's dose-response due to baseline growth rates. For each compound dose-series and strain, GR < 0 was truncated to 0, GR > 1 was compressed to be close to 1 (while maintaining rank order of the strains), and two dummy/null measurements of GR = 1 at 1/8 and 1/16 (3 and 4 dilutions below) the lowest assayed dose were concatenated to the observed GR measurements in order to aid in curve-fitting and more strictly constrain GR between 0 and 1.

$$GR_{compressed\ for\ curve-fitting} = f(GR) = \begin{cases} 0, & GR < 0 \\ GR, & 0 \le GR \le 1 \\ 1 + \dfrac{GR - 1}{10}, & GR > 1 \end{cases}$$

The null datapoints (at concentrations very close to 0 uM, i.e., vehicle-like) incentivized the upper bound/asymptote on the dose-response model fits for each strain to be GR = 1 (uninhibited, vehicle-like growth) which is the simple and expected biological result had the compound been screened at a complete and infinitesimally dilute dose-series. Importantly, for strains that were significantly inhibited with GR < 1 across the entire dose-series of a compound, the use of the null datapoints aided in achieving monotonically decreasing functions that estimated return to uninhibited growth with GR = 1 at some rate as compound concentration decreased unlike unconstrained best-fits, which could be monotonically increasing or nonmonotonic functions that are less biologically applicable and the result of overfitting to a limited dose-series. The null datapoints were included for dose-response curve fitting and otherwise excluded from all other analysis. 5-parameter logistic, 4-parameter logistic, and Gompertz function best-fit curves were found and the function with the lowest Akaike information criterion (AIC) was selected[14]. The values of the selected function at the compound's tested doses were taken as the strain's curve-fit, de-noised GR growth inhibition values and combined with all other strains to make up a curve-fit GR CGI profile for each assayed concentration of the compound.

Strain curve-fit GR, with the benefits of reducing dose-to-dose noise in the assay and stricter constraints on GR to be between 0 and 1, was then used to estimate compound activity as the lowest strain curve-fit GR achieved at the highest tested dose of the compound. Each condition's activity more generally was estimated as the lowest strain curve-fit GR achieved at each tested dose. Compounds that inhibited at least one strain significantly enough to achieve a curve-fit GR ≤ 0.3 at the highest tested dose (i.e., achieving a strain $GR_{70}$ following Hafner et al.) were defined as active and all others that didn't meet this criterion were considered inactive (or minimally active) in the assay[3]. Using raw, quantile normalized GR and description of compound activity as the lowest GR achieved over any tested dose did not significantly change the proportion of compounds identified as active or inactive or the overall finding of higher MOA prediction sensitivity and precision for active compounds.

### Supplementary Note 7 | Unsupervised clustering of CGI profiles from each MOA in the reference set

To cluster sGR CGI profiles from each MOA category, we applied spectral clustering in Matlab, a robust technique that has been shown to segregate and cluster complex data more accurately than other common methods[15,16]. This process involved four steps. First, we calculated Pearson correlation as the similarity metric between condition sGR profiles. Second, we constructed a nearest-neighbor graph for each MOA class, connecting conditions to others within the same MOA that were within the top 20 closest conditions across all of the reference set (**Extended Data Fig. 4**). For each condition *i*, the correlation coefficients between it and all other conditions in the reference set were ranked such that rank 1 and rank 9,427 were respectively the highest correlated/most similar and lowest correlated/least similar conditions to condition *i* across all reference CGI profiles. For every pair of conditions *i* and *j*, their ranks in Pearson correlation were averaged to symmetrize the similarity matrix and if their average rank in Pearson correlation was ≤ 20 then the pair of conditions were connected as nearest neighbors. This is stored as an *n*-by-*n* adjacency matrix *W* where *n* is the number of conditions (CGI profiles) in the MOA and entries between connected conditions are 1 and disconnected conditions are 0. Third, we estimated the number of clusters *k* in each MOA by representing its similarity graph as a graph Laplacian and using an eigengap heuristic. The normalized symmetric Laplacian matrix (Ng-Jordan-Weiss) was calculated as $L_s=D_g^{-1/2}LD_g^{-1/2}$ where $L=D_g-W$ and $D_g$ is the degree matrix of *W*, an *n*-by-*n* diagonal matrix with entry $D_g(i,i)$ equal to the total number of connected conditions in the MOA to condition *i* (i.e., the sum of row *i* in the adjacency matrix *W*)[15]. The eigenvectors and eigenvalues of the Laplacian matrix $L_s$ were computed and the number of $k$ clusters in the MOA were estimated as the average (rounded up to the nearest integer) of $k_{zero\_plus\_one}$ and $k_{gap}$ where $k_{zero\_plus\_one}$ is the index of the smallest non-zero eigenvalue and $k_{gap}$ is the eigenvalue where the gap or change in consecutive eigenvalues was largest. While, in theory, the number of eigenvalues equal to zero $\mathrm{k_{zero}}$ indicates the number of connected components or clusters for perfectly separable data, real-world data is noisy and $k_{zero\_plus\_one}$ and $k_{gap}$ have each been proposed as appropriate estimates for the number of connected components and termed spectral gap or eigengap heuristics[15,16]. We found that estimating $k$ as the average of $k_{zero\_plus\_one}$ and $k_{gap}$ resulted in the greatest overall performance in our method. We reasoned that this could be due to $k_{zero\_plus\_one}$ underfitting for some MOAs resulting in too few and large of clusters and $k_{gap}$ overfitting for some MOAs resulting in too many and small of clusters. Ultimately, this allowed for estimation of the number of $k$ clusters in each MOA to be fully automated and standardized. Finally, using the spectralcluster Matlab function, for each MOA and its estimated $k$, the $k$ clusters were found by reducing the condition space ($n$-by-$n$) to a lower-dimensional ($n$-by-$k$) space using the first $k$ eigenvectors of the Laplacian matrix and assigning each condition $i$ in $n$ to one of $k$ clusters using k-means clustering[17,18]. Specifically, the first $k$ eigenvectors of $L_s$ are combined column-wise into an $n$-by-$k$ matrix $V$ and each row of $V$ is normalized to have unit length. The $n$ points (conditions) are then clustered in the embedded $k$-dimensional feature space using k-means clustering and assigned into one of $k$ clusters. This process was repeated for each MOA separately and the cluster identifiers and CGI profiles members of each cluster were stored in GMT file format. For reproducibility, the default Matlab random number generator (i.e., Mersenne Twister generator with seed 0) was initialized using the rng function, the maximum number of computational threads was set to 4 using the maxNumCompThreads function, and the order of conditions in the W, $\mathrm{D_g}$, L, and $\mathrm{L_s}$ matrices was sorted by hierarchical clustering of the Pearson correlation matrix using CmapM's hclust function for each MOA's iteration of spectral clustering.

### Supplementary Note 8 | PCL cluster definition and similarity score

We defined and calculated a PCL similarity score between each test CGI profile and each cluster from unsupervised, spectral clustering as the median of the Pearson correlation coefficients between the test CGI profile and all reference set CGI profiles within the cluster. Clusters whose highest similarity scoring, reference set CGI profile shared the same annotated MOA as the cluster members (i.e., one of the cluster CGI profiles themselves or a CGI profile from a compound in the same MOA but was a member of a different cluster) were kept for use in making MOA predictions and defined as PCLs. However, clusters whose highest similarity scoring reference set CGI profile was from a compound of a different annotated MOA than the cluster members were discarded as "non-predictive" clusters whose pattern of strain inhibition was non-specific to one single MOA and therefore were difficult to interpret for predicting MOA on unknown compounds. When calculating PCL similarity score between a reference set CGI profile and a cluster it was a member of, self-similarity (Pearson correlation of 1) was excluded from the median calculation. Including self-similarity in this calculation would unfairly bias higher similarity scores for CGI profiles in the cluster than CGI profiles from new, unknown compounds whose cluster membership is unknown (had their MOAs been annotated and included in the reference set) and reference set CGI profiles that belong to other clusters and would otherwise bias comparison of similarity scores when defining the thresholds to be used for making high-confidence MOA predictions. PCL similarity scores for each CGI profile to each PCL were stored in GCTx annotated data matrix format.

### Supplementary Note 9 | One-vs-rest (OvR) binary classification strategy

We posed MOA prediction on unknown compounds as a multi-class classification problem using a one-vs-rest binary classification strategy in which a single classifier per PCL is trained with reference set compounds of its MOA as positive samples and reference set compounds of all other MOAs as negative samples; when applied to an unknown compound it outputs a real-valued confidence score of the unknown compound acting via the same MOA as the PCL[19-21]. For each PCL, we had the set $S = \{(x_1, y_1), \ldots, (x_{437}, y_{437})\}$ of 437 reference set compound training samples where $x_i \in X$ was the maximum similarity score of the *i*th compound (over all of its CGI profiles, i.e, doses) to the PCL, $y_i \in Y$ is 1 if the compound shares the same MOA as the PCL and 0 otherwise. Each PCL's classifier is a function that maps a similarity score x to a confidence score (ranging from 0 to 1) estimating the likelihood of the compound acting via the PCL's annotated MOA. Specifically, it is the proportion of reference set compounds with similarity score ≥ x that share the PCL's MOA, i.e., the precision (or positive predictive value, i.e. PPV) of MOA assignment if the similarity score x was used as the threshold above which all reference set compounds are labeled 1 and all reference set compounds below are labeled 0. Confidence scores (i.e., PPV) at each $x_1, \ldots, x_{437}$ were computed using *perfcurve* in Matlab. While the initial mapping was calculated using the highest similarity scores from each of the 437 compounds to avoid repeated tallying of non-independent samples (multiple CGI profiles of the same compound), the mapping was extended to similarity scores from all reference set CGI profiles via next neighbor interpolation (*interp1* in Matlab) – the confidence score for a similarity score from an *unobserved* CGI profile is the confidence score of the next highest similarity score from an *observed* CGI profile (i.e., the most similar profile for a reference set compound). The rationale follows: for an *unobserved* CGI profile from reference set compound *i* with similarity score $\bar{x}_i < x_i$ and lowest *observed* similarity score $x > \bar{x}_i$, replacement of $(\bar{x}_i, y_i)$ for $(x_i, y_i)$ in set S (e.g., the dose with similarity score $x_i$ wasn't screened and $\bar{x}_i$ was instead the maximum similarity score of compound *i* to the PCL) during initial mapping would compute a confidence score for $\bar{x}_i$ equal to the confidence score for x. In other words, the proportion of reference set compounds with similarity scores $\geq \bar{x}_i$ excluding $x_i$ is the same as the proportion with similarity scores ≥ x including $x_i$. We reasoned that compounds would be the most appropriate, independent data points to tally as evidence supporting an MOA call for a PCL, as opposed to doses which could be from a single compound not shared across the MOA and lead to strong biases toward large MOAs (where the relative differences in the number of CGI profiles per MOA is ≈10 times greater than the number of compounds). The full mapping of similarity score to confidence score for every reference set CGI profile to every PCL was stored and referred to as the full model. High-confidence similarity score thresholds were defined for each PCL as the lowest similarity score where confidence score = 1.

### Supplementary Note 10 | Applying PCL classifiers to CGI profiles from unknown compounds

When applying a PCL classifier to an unknown compound, the similarity scores of its CGI profiles were mapped to confidence scores via linear interpolation (*interp1* in Matlab) using the mapping of similarity score to confidence score trained on the reference set. This approach considers the distance (or difference) to an unknown CGI profile's similarity score x and the two reference set CGI profile similarity scores it is between $x_n > x > x_m$ when estimating the confidence score for x based on the confidence scores of $x_n$ and $x_m$. Confidence scores were estimated for every unknown CGI profile to every PCL and stored in GCTx annotated data matrix format.

### Supplementary Note 11 | PCL-based MOA predictions for unknown compounds

For each unknown compound, its most likely MOA in the reference set was predicted as the PCL with the highest confidence score over one or more of its CGI profiles, following the typical one-versus-rest multi-class classification strategy[19-21]. Since there can be PCL classifiers (of the same or different MOAs) with equally high confidence scores for an unknown compound, we made one MOA prediction for every compound via a majority rules tiebreaker in which the PCL that had the highest confidence score over the greatest number of doses (CGI profiles) and highest similarity score overall, if necessary, was selected. The predicted MOA was considered a high-confidence MOA assignment if the PCL confidence score was equal to 1. A compound's MOA was considered "uncertain" if its highest PCL confidence score was below 1. For unknown compounds lacking a high-confidence MOA assignment, their predicted MOAs and the strength of their PCL confidence scores could still be considered, to provide mechanistic insight for more unknown compounds with slightly reduced confidence (**Extended Data Fig. 9**).

The above approach can be taken to separately predict MOA at every data level for a compound – dose (i.e., CGI profile), specific lot or batch of a compound in a particular screening wave (i.e., proj_broad_id identifier), specific lot or batch of a compound over all screening waves (i.e., broad_id identifier), or compound (i.e., pert_id identifier) – for comparing behavior between screens or commercial and synthetic batches of a compound (**Extended Data Table 3**, **Supplementary Data 4**). As well, for compounds with equally high confidence scores to PCLs from multiple MOAs, each MOA and the reference compounds from each of the PCLs can be considered holistically (i.e., multi-label classification) and motivate follow-up experimental validation and rule-out of each of the MOAs (**Supplementary Data 5**).

### Supplementary Note 12 | Leave-one-out cross validation and tested PCL analysis variations

For each reference set compound, its CGI profiles were removed from the reference set and all CGI profiles from the remaining 436 reference set compounds were used for unsupervised clustering, PCL cluster definition and similarity scoring, and for training PCL classifiers to map similarity score to confidence score as described above. Using the mappings of similarity score to confidence score for each PCL, confidence scores for each of the left-out reference set compound's CGI profiles were estimated via linear interpolation, as described above for unknown compounds using the full model. The estimated confidence scores of each reference set compound to each PCL from its iteration of LOOCV were combined. A single MOA prediction was made for each reference set compound based on the highest confidence scoring PCL (using the same majority rules tiebreaker as above) and considered to be a high-confidence MOA assignment if the confidence score was 1. True positives and false positives in LOOCV were tallied as the number of reference set compounds whose high-confidence MOA assignments were concordant or discordant, respectively, with their annotated MOAs and compounds that were "uncertain" (i.e., all PCL confidence scores < 1) were tallied as false negatives since their annotated MOAs were unable to be predicted with high-confidence. Micro-averaged precision, sensitivity, and F1 score statistics were calculated using the counts of true positives, false positives, and false negatives in LOOCV[22] with specific focus on reference set compounds that were identified as active in the screen (**Extended Data Table 1**). 19 compounds were the sole representative of their annotated MOAs in the reference set and cannot be correctly predicted in LOOCV – since no PCLs of their MOA or other data annotated to their MOA exist in the training set when they are withheld.

To evaluate the importance of the unsupervised clustering step in identifying highly similar clusters of CGI profiles within each MOA and filtering out less informative CGI profiles, we repeated the standard LOOCV procedure described above but, in each iteration, took the complete set of CGI profiles from each MOA as one comparison group per MOA instead of applying spectral clustering. The following steps of PCL cluster definition and similarity scoring, PCL classifier training, estimation of PCL confidence scores and MOA prediction for each left-out reference set compound, and calculation of micro-average accuracy statistics remained the same with up to 71 groups (one for each MOA) as the starting point instead of the clusters from spectral clustering (1,117 clusters on average) (**Extended Data Table 1**).

To evaluate the importance of the PCL classifier training step and defining high-confidence similarity score thresholds unique to each PCL using confidence score, we tested the standard LOOCV procedure described above but each left-out reference set compound's MOA prediction was made to the PCL with the highest similarity score over any of its CGI profiles. Balancing precision and sensitivity equally, ROC analysis was performed in R, using the set of PCL similarity scores from each of the reference set compounds and binary labels (1 = agree, 0 = disagree) if the PCL matched the annotated MOA of the compound, to find the optimal similarity score threshold above which a known or unknown compound would be assigned the PCL's MOA with "high-confidence" or otherwise considered "uncertain"[23]. The similarity score thresholds found in this approach were more stringent than the high-confidence similarity score thresholds found and defined for most PCLs on an individualized basis (**Extended Data Table 1**, **Extended Data Fig. 7, 8**). Compounds were assigned an MOA if their similarity score surpassed the screen-wide similarity score threshold found in the ROC analysis (e.g., PCL similarity score ≥ 0.649 found for reference set compounds from multi-compound MOAs), otherwise they were considered "uncertain". The preceding steps of unsupervised clustering and PCL cluster definition and similarity scoring and the following step to calculate micro-average accuracy statistics were the same as the standard LOOCV procedure.

To further evaluate the benefit of the PCL analysis method and the corroboration of multiple similar reference set CGI profiles (PCL clusters) when comparing to test compounds, we evaluated the predictive performance of a simple 1-nearest neighbor approach where the MOA assigned to a test compound is the MOA of the reference set CGI profile with the highest Pearson correlation coefficient in LOOCV. For each left-out reference set compound, all its CGI profiles were removed from the reference set and its 1-nearest-neighbor (over any of the left-out compound's screened compound-doses) was selected as the CGI profile from the remaining reference set with the highest Pearson correlation coefficient. The MOA of each compound's 1-nearest neighbor in LOOCV was compared to their annotated MOA to calculate the accuracy of the predictions. Micro-averaged precision, sensitivity, and F1 score statistics were calculated using the counts of true positives and false positives in LOOCV with specific focus on reference set compounds that were identified as active in the screen (**Extended Data Table 1**)

### Supplementary Note 13 | Tanimoto similarity calculation

The pairwise Tanimoto similarity coefficient was calculated between each compound in the GSK set and BRD4310 and each compound in the reference set using the RDKit library (v.2020.09.1) in Python (v.3.7.12). For each compound, its SMILES was converted to an RDKit molecular object and its Morgan fingerprint was computed using a neighborhood size of 2 and all other default parameters[24]. The Tanimoto similarity coefficient was calculated for every pair of test and reference set molecular fingerprints. For each GSK compound and BRD4310, the reference compound with the highest Tanimoto similarity coefficient was taken as its nearest chemical neighbor in the reference set (**Extended Data Table 2**, **Supplementary Data 6**).

### Supplementary Note 14 | Chemical synthesis and characterization

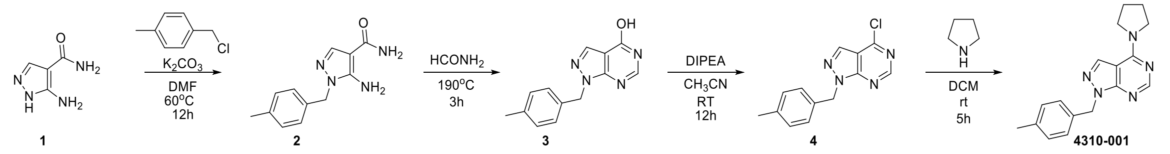


Synthesis of 4-(1-pyrrolidinyl)-1-[(p-tolyl)methyl]-1,2,5,7-tetraaza-1H-indene (**BRD4310**, i.e., **4310-001**):

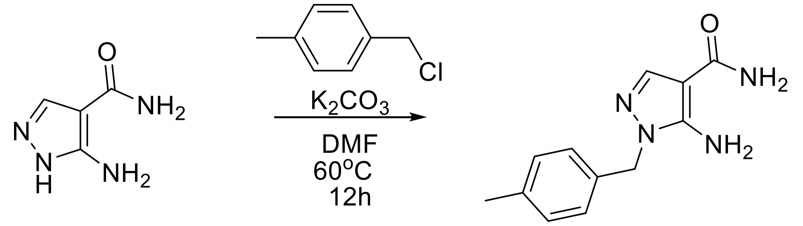


**1** **2**

To a mixture of 5-amino-1H-pyrazole-4-carboxamide (5 g, 39.68 mmol, 1 eq) and $K_2CO_3$(10.95g, 79.37 mmol, 2 eq) in DMF (dimethylformamide, 198 mL) was added 1-(chloromethyl)-4-methylbenzene (11.19g, 39.68 mmol, 2 eq). The resulting mixture was stirred at 60°C for 12 h. After being cooled to room temperature, the reaction mixture was diluted with water (50 mL) and extracted with EA (ethyl acetate, 50 mL×3). The combined organic phase was dried over anhydrous $Na_2SO_4$ and concentrated in vacuo. The residue was purified by silica gel chromatography to give 2.8 g of 5-amino-1-(4-methylbenzyl)-1H-pyrazole-4-carboxamide. Yield was 30%. LC-MS: $[M+H]^+$ = 231.4. $t_R$ = 1.68.

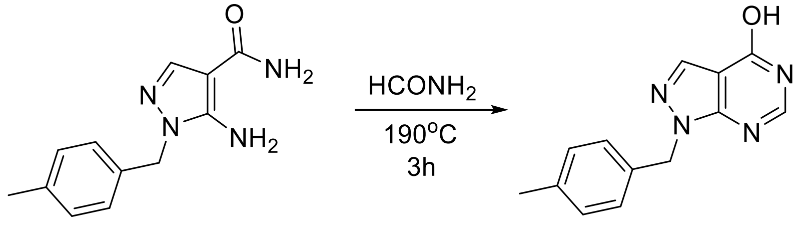


2 3

A mixture of 5-amino-1-(4-methylbenzyl)-1H-pyrazole-4-carboxamide (3 g, 13 mmol, 1 eq) and formamide (26 mL) was stirred at 190°C for 3 h. After being cooled to room temperature, the reaction mixture was diluted with DCM (dichloromethane, 50 mL), washed with water (50 mL×3), dried over anhydrous $Na_2SO_4$ and concentrated in vacuo. The residue was purified by silica gel chromatography to give 2.3 g of 1-(4-methylbenzyl)-1H-pyrazolo[3,4-d]pyrimidin-4-ol. Yield was 73%. LC-MS: $[M+H]^+$ = 241.1. $t_R$ = 1.71.

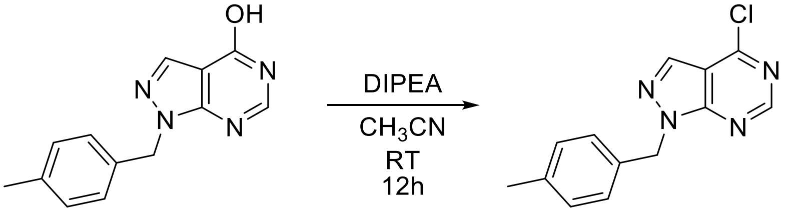


3 4

To a mixture of 1-(4-methylbenzyl)-1H-pyrazolo[3,4-d]pyrimidin-4-ol (1 g, 4.16 mmol, 1 eq) in $CH_3CN$ (14 mL) were added $POCl_3$ (5.74 g, 37.5 mmol, 9 eq) and DIPEA (N,N-Diisopropylethylamine, 1.18g, 9.15 mmol, 2.2 eq). The resulting mixture was stirred at 0°C for 15 min and at room temperature for 12 h. The reaction mixture was diluted with $CH_3CN$ (10 mL) and quenched with $H_2O$ (10 mL). The resulting mxiture was adjusted with $NaHCO_3$ to pH = 8. The mixture was extracted with DCM (dichloromethane, 100 mL). The extracts were washed with brine, dried over anhydrous $Na_2SO_4$ and concentrated in vacuo. The residue was purified by silica gel chromatography to give 900 mg of 4-chloro-1-(4-methylbenzyl)-1H-pyrazolo[3,4-d]pyrimidine. Yield was 83% LC-MS: $[M+H]^+$ = 259.1, $t_R$ = 1.71.

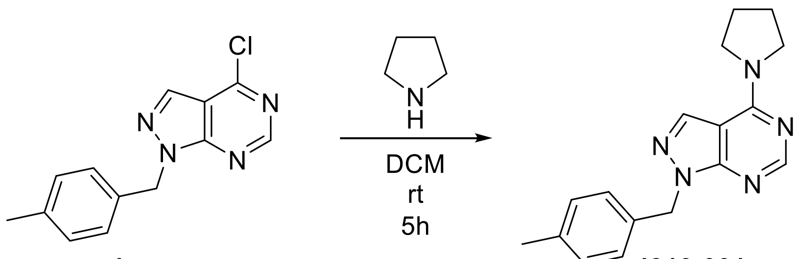


**4** **4310-001**

To a mixture of 4-chloro-1-(4-methylbenzyl)-1H-pyrazolo[3,4-d]pyrimidine (300 mg, 1.16 mmol, 1 eq) in DCM (dichloromethane, 4 mL) was added pyrrolidine (1.94 mL, 23.25 mmol, 20 eq). The resulting mixture was stirred at room temperature for 5 h. The reaction mixture was diluted with water (20 mL) and extracted with EA (ethyl acetate, 30 mL×3). The combined organic phase was dried over anhydrous $Na_2SO_4$ and concentrated in vacuo. The residue was purified via prep-HPLC to provide **BRD4310** (i.e., **4310-001**) in 20% yield (68 mg). LC-MS: $[M+H]^+$ = 295.4. $t_R$ = 1.86. $^1H$ NMR (400 MHz, DMSO-$d_6$) δ 8.37 (d, J = 9.9 Hz, 1H), 8.31 (s, 1H), 7.11 (d, J = 8.7 Hz, 4H), 5.50 (s, 2H), 3.84 (t, J = 6.7 Hz, 2H), 3.68 (t, J = 6.7 Hz, 2H), 2.25 (s, 3H), 2.09 (dt, J = 13.0, 6.5 Hz, 2H), 1.99 (dd, J = 13.1, 6.5 Hz, 2H).

**BRD4310 $^1H$ NMR and LC-MS:**

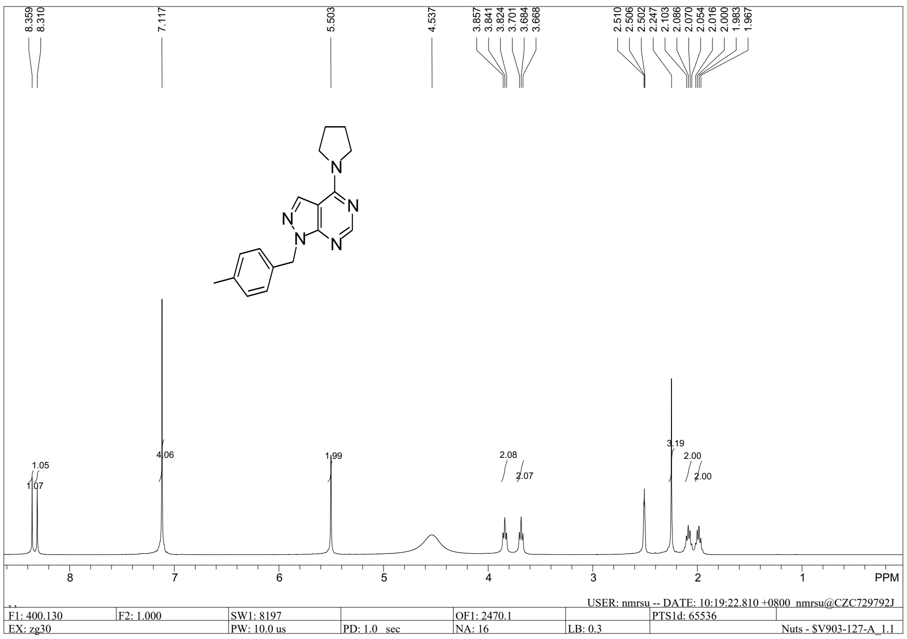

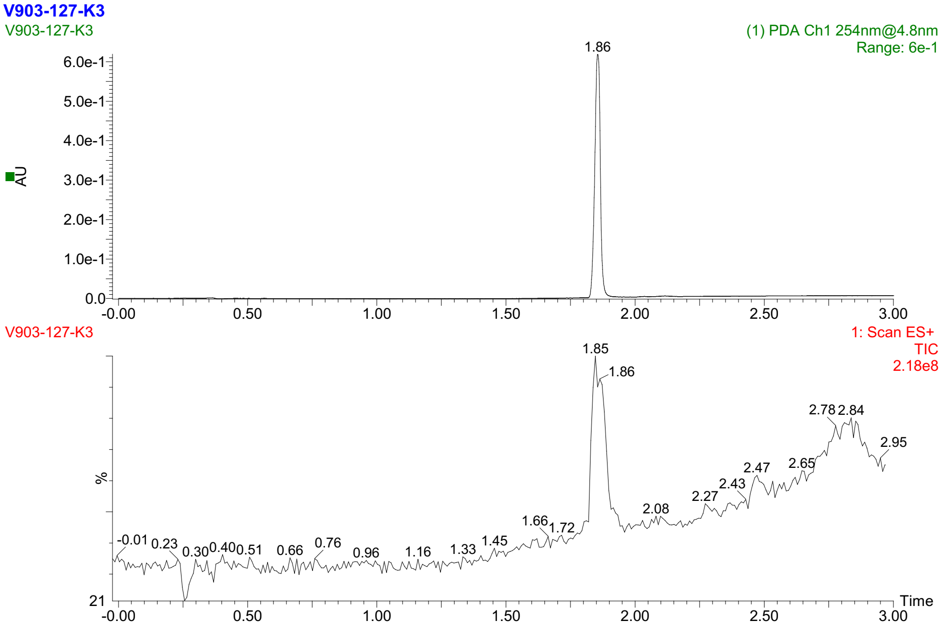


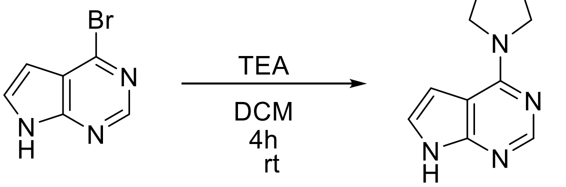


**1** **IntermediateB**

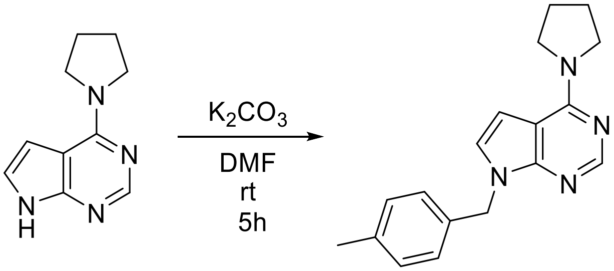


**Intermediate B** **4310-008**

Synthesis of 4-(1-pyrrolidinyl)-1-[(p-tolyl)methyl]-1,5,7-triaza-1H-indene (**BRD4310a**, i.e., **4310-008**):

To a mixture of 4-bromo-7H-pyrrolo[2,3-d]pyrimidine (5 g, 25.3 mmol, 1 eq) and TEA (triethylamine, 5.1 g, 50.6 mmol, 2 eq) in DCM (dichloromethane, 100 mL) was added pyrrolidine (2.15g, 30.3 mmol, 1.2 eq). The resulting mixture was stirred at room temperature for 4 h. The reaction mixture was diluted with water (100 mL). The organic phase was separated and the aqueous phase was extracted with EA (ethyl acetate, 100 mL×3). The combined organic phase was washed with brined, dried over anhydrous $Na_2SO_4$ and concentrated in vacuo. The residue was purified by silica gel chromatography to give 4 g of **Intermediate B**. Yield was 84%. LC-MS: $[M+H]^+$ = 189.1.

To a mixture of **Intermediate B** (100 mg, 0.53 mmol, 1 eq) in DMF (dimethylformamide, 2.6 mL) were added 1-(chloromethyl)-4-methylbenzene (150 mg, 1.06 mmol, 2 eq) and $K_2CO_3$ (146 mg, 1.06 mmol, 2 eq). The resulting mixture was stirred at room temperature for 5 h. The reaction mixture was filtered and the filtrate was concentrated in vacuo. The residue was purified via prep-HPLC to provide **BRD4310a** (i.e., **4310-008**) in yield 41% (86 mg). LC-MS:$[M+H]^+$ = 293.1, $t_R$ = 1.8min. $^1H$ NMR (400 MHz, $CDCl_3$) δ 8.44 (s, J = 10.1 Hz, 1H), 7.13 (dd, J = 8.6, 2.7 Hz, 3H), 7.10 (d, J = 3.5 Hz, 1H), 6.72 (t, J = 6.9 Hz, 1H), 5.38 (s, 2H), 3.97 (s, 4H), 2.32 (s, 3H), 2.16 (s, 4H).

**BRD4310a $^1H$ NMR and LC-MS:**

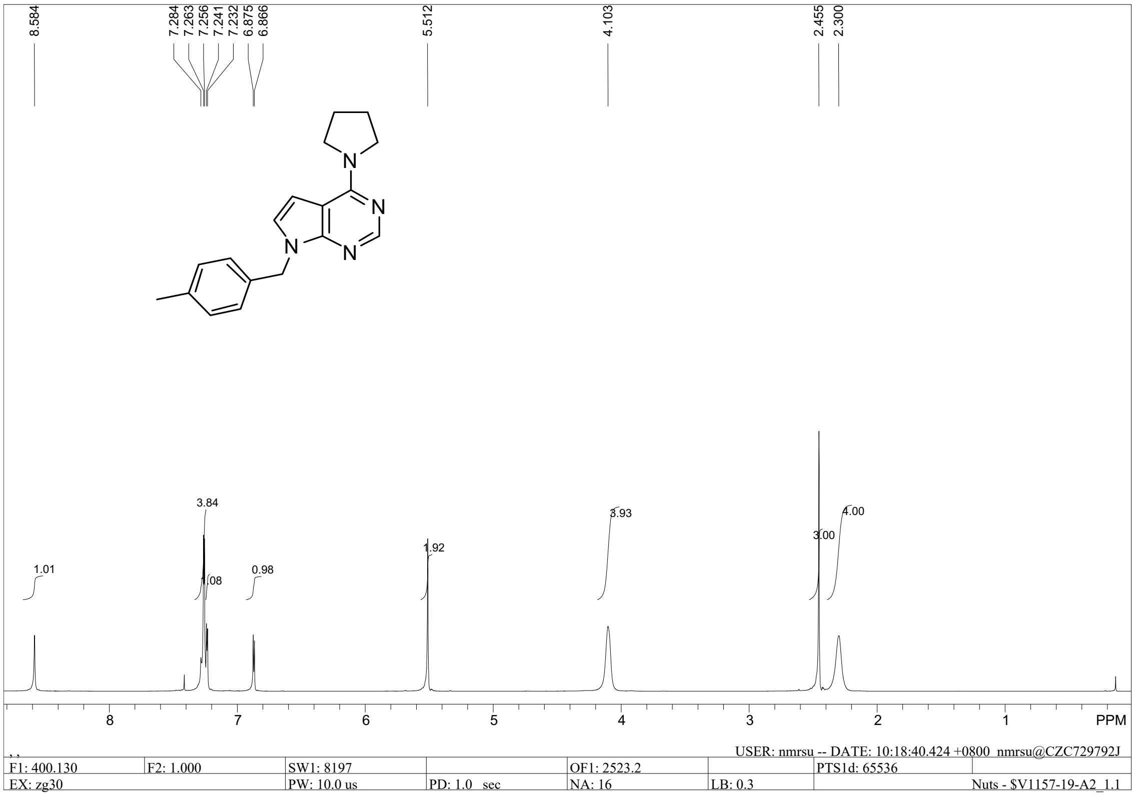

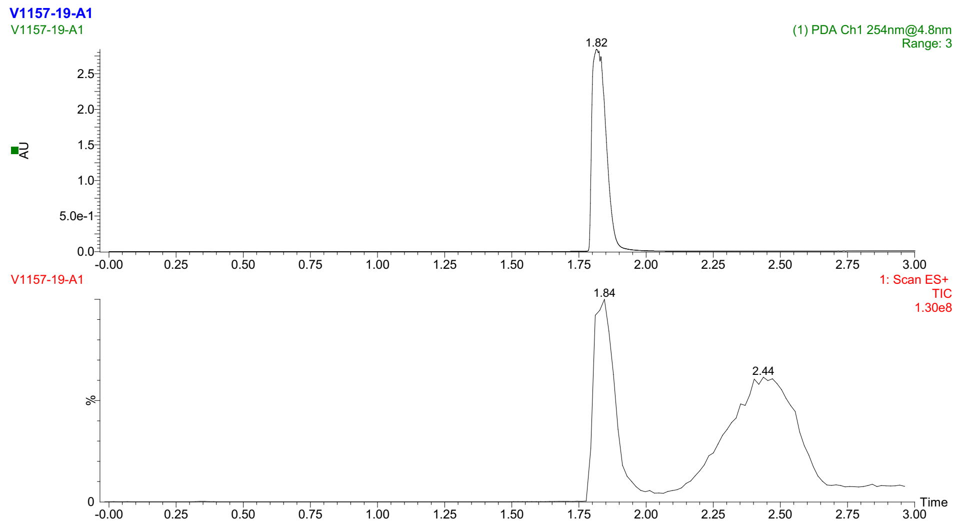
V1157-19-A1
V1157-19-A1
(1) PDA Ch1 254nm@4.8nm
Range: 3
1.82
AU
2.5
2.0
1.5
1.0
5.0e-1
0.0
-0.00 0.25 0.50 0.75 1.00 1.25 1.50 1.75 2.00 2.25 2.50 2.75 3.00
V1157-19-A1
1: Scan ES+
TIC
1.30e8
1.84
2.44
%
0
Time
-0.00 0.25 0.50 0.75 1.00 1.25 1.50 1.75 2.00 2.25 2.50 2.75 3.00
